# The Vizitig pangenome and pantranscriptome explorer

**DOI:** 10.1101/2025.04.19.649656

**Authors:** Bastien Degardins, Charles Paperman, Camille Marchet

## Abstract

Vizitig is the first platform for real-time exploration and querying of DNA and RNA sequence de Bruijn graphs across many samples, unifying visualization, metadata, and flexible search. It constructs compacted colored de Bruijn graphs from raw sequencing reads and reference sequences, then provides an interactive web interface for graph exploration. It integrates raw and reference-based data, handles complex variation, and provides a human-readable feature-based (also referred to as metadata) query language with scalable graph loading. Its domain-specific query language supports composable searches combining sequences of arbitrary size, genomic features such as gene or exon identifiers, and experimental factors such as sample identifier or abundance thresholds. On-demand subgraph loading retrieves only regions of interest, enabling interactive exploration of large datasets in the graphical user interface without loading the entire graph into memory. We demonstrate Vizitig’s capabilities through case studies in pantranscriptomics and pangenomics. In pantranscriptomics, we recover fusion transcript breakpoints on long and short reads. In pangenomics, we explore sequence variations across yeast, rice, nematode, and human pangenomes. Vizitig scales from small virus genomes to human-scale pangenomes (our largest experiment comprises up to 20 assembled human haplotypes, on a laptop). Vizitig enables fast, reproducible analysis in both pangenomics and pantranscriptomics while providing a deployable and user-friendly working environment.

## Introduction

Advances in high-throughput sequencing allow the construction of large genome and transcriptome collections, driving discoveries in evolution, disease, and functional biology. Biological variants often require comparisons across several individuals, strains, or samples. Tools such as BLAST and IGV (Thorvaldsdóttir et al. 2013) have highlighted the power of interactive exploration, but remain tied to linear reference models. We now need models that integrate diverse sequence collections, such as population-scale pangenomes (Holley et al. 2026; Gao et al. 2023; Leonard et al. 2024) and multi-sample transcriptomes (pantranscriptomes).

Graph-based representations have emerged as a natural framework for encoding the diversity present in multi-sample datasets such as pangenomes. Variation graphs, built from whole-genome alignments, capture structural variants and rearrangements relative to either a reference coordinate system or relative to the samples themselves by doing all-versus-all alignment. Dedicated builders such as Minigraph (Li et al. 2020), Minigraph-Cactus (Hickey et al. 2024), and PGGB (Garrison et al. 2024) have matured considerably, culminating in the construction of the first draft human pangenome reference (Liao et al. 2023). De Bruijn graphs offer an alignment-free alternative: they decompose input sequences into overlapping *k*-mers and represent them as nodes connected by suffix– prefix overlaps, capturing all variation, including single nucleotide changes, as topological features of the graph. Colored extensions of the de Bruijn graph further associate each node with the set of samples in which it occurs, enabling comparative analyses without a prior alignment step (Marchet et al. 2021). Formally, the main difference lies in the definition of the graphs. A variation graph is easy to interpret for a human but is not formally defined, whereas de Bruijn graphs are well-defined but can be hard to interpret biologically speaking. Vizitig aims at bridging this gap. Current pangenomes rely on graphs whose topology can vary across methods and software versions (Chikhi et al. 2024).

Existing software to interact with pangenomes offers valuable yet domain-specific functionality: assembly (Wick et al. 2015) and variation graph viewers, toolkits (Guarracino et al. 2022), comparative genomics and phylogenetics frameworks (Jonkheer et al. 2022). Visualization tools such as Bandage (Wick et al. 2015), Sequence Tube Map (Beyer et al. 2019) and VRPG (Miao and Yue 2025) each address graph rendering, but none integrates construction, annotation, querying, and visualization within a single pipeline. The odgi toolkit (Guarracino et al. 2022) addresses graph manipulation and visualization, but not graph construction.

Conversely, pantranscriptomes still lack a factorized, graph-based representation. Although splicing graphs and transcript-level quantification tools exist, no current platform allows interactive, multi-sample exploration of RNA-seq data in a graph context while simultaneously supporting feature-driven queries and sequence extraction. Yet such exploration is increasingly needed: detecting rare events such as fusion transcripts or circular RNAs requires inspecting graph neighborhoods that span non-canonical junctions absent from standard annotations.

In both pangenomic and pantranscriptomic contexts, a key challenge is to abstract the complexity arising from many accumulated variants, most irrelevant to a given question, while still enabling precise sequence extraction from regions or individuals of interest.

Here, we present Vizitig, a tool that integrates pangenome or pantranscriptome graph construction and exploration, offering query-driven access to DNA/RNA datasets through either a graphical web interface or a command line interface. Our aims were (i) to build, annotate, query, and visualize colored de Bruijn graphs from raw reads and assemblies within a single tool; (ii) to make these graphs accessible to non-specialists through a composable, feature-aware query language and on-demand subgraph loading; and (iii) to show, through pantranscriptomic and pangenomic case studies ranging from yeast to human, that this approach recovers biologically relevant events and scales to large datasets on a laptop.

## Results

### Vizitig overview

Rather than aligning sequences to a fixed reference coordinate system, Vizitig represents each dataset in a graph built directly from short overlapping subsequences (*k*-mers), an alignment-free strategy that scales to large, multi-sample collections and captures variation as branch points rather than as deviations from a single reference. This corresponds to the colored de Bruijn graph model (Marchet et al. 2021). Vizitig constructs compacted colored graphs of this kind from input sequences, and provides an interactive interface and a permissive query language for their exploration (Figure 1), using principles shared with assemblies and large-scale indexes (Karasikov et al. 2025). More precisely, *k*-mers are first mapped to node ids in the graph during the indexing phase, enabling fast *k*-mer lookup to improve query time. Second, a database links node ids with features, sample occurrences, and abundances. This tight coupling between graph topology and metadata eliminates the need to switch between separate tools for querying and visualization, a workflow that currently limits the adoption of graph-based analyses by non-specialists. A logic layer supports expressive, composable queries (union, intersection, difference) across three modalities:

**Figure 1:**
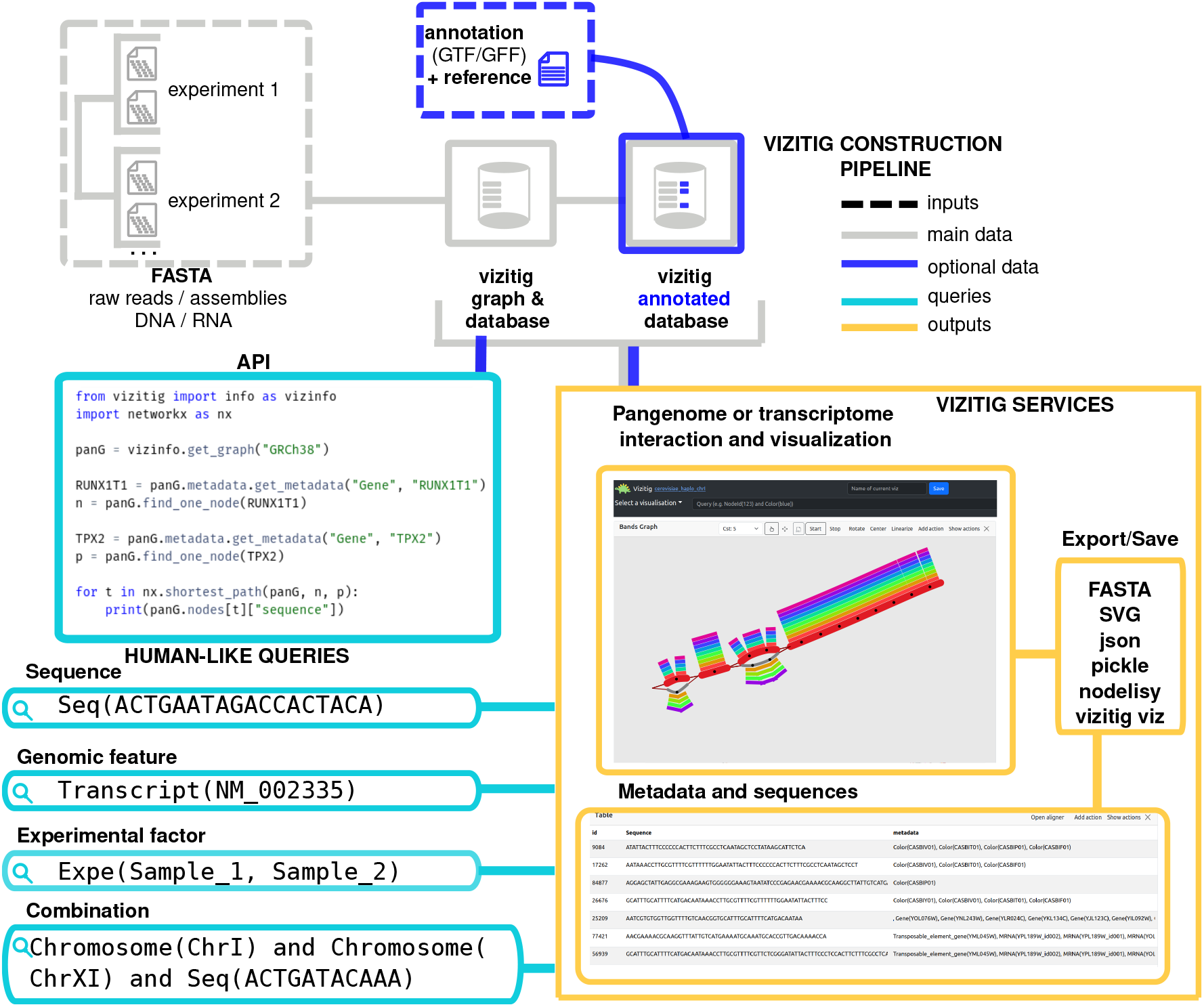
Overview of the Vizitig explorer tool. Vizitig integrates diverse data types (reads and references, DNA and RNA) in a de Bruijn graph. Users can filter and query graphs by metadata and sequence traits. An interactive web interface streams only relevant subgraphs, enabling scalability. Results, metadata, and views can be exported for sharing or downstream analysis.

1. Sequence of arbitrary span with abundance conditions (regardless of the size of *k*);
2. Features (e.g., transcript ID, gene name, chromosome);
3. Experimental factors (e.g., strain, sample).

To our knowledge, Vizitig is the first method to allow this set of queries and a combination thereof, e.g., “retrieve a given mutant *BRCA1* sequence in two selected samples with abundance ≥10”. Bandage (Wick et al. 2015) also allows sequence queries, but relies on BLAST (Altschul et al. 1990) to do so. The relational model behind the database allows this higher level of compositionality, which is currently possible in no other tool, as well as an efficient query execution plan. Classic variation-graph-based renderers, which are limited to sequence, coordinate, or node id queries, lack such compositionality and optimizations due to the linearity of GFA formats.

Another novelty is dynamic, on-demand subgraph loading: Vizitig retrieves only the regions matching a query and lets users selectively expand the visible neighborhood using the same query logic (e.g., expand 100 nodes that appear in sample X, starting from the current subgraph). At any point during exploration, users can inspect the sequence, features, sample occurrences, and abundance of any node, a level of interactivity not available in other graph rendering tools. A large set of visual transformations can be applied to the loaded subgraph (e.g., coloring the nodes with their origin samples, highlighting variants).

Vizitig integrates pangenome and pantranscriptome construction, querying, and interactive exploration within a single tool (Supplemental Figs. S1–S2; Supplemental Tables S1–S3, S6), a combination we show is not offered by existing solutions. Beyond this integration, Vizitig is fast: parsing, indexing, and querying scale to large datasets, with graph construction scaling linearly and query responses returned in seconds (Supplemental Tables S1–S2, S5). In the following we illustrate these capabilities through several case studies spanning pantranscriptomics and pangenomics.

## Detection of rare transcriptomic events

Our first case study concerns transcriptomic events relevant to cancer, typically rare or weakly expressed, and therefore difficult to detect: fusion transcripts (exons joined from different genes) and circular RNAs (covalently linked 5′–3′ loops that linear models capture poorly). For each, we selected known positives from prior studies and constructed Vizitig de Bruijn graphs using the corresponding samples and RefSeq transcripts. Datasets are listed in Supplemental Table S1, and all graphs, visualizations, and command lines are released for reproducibility (see the Zenodo repository in Methods).

We re-analyzed RNA-seq data containing two circular RNAs, *MAN1A2* and *FBXW4*, originally described by Mercer et al. (2015) (Figure 2A). Starting from querying gene-IDs, we could visually detect alternative back-splicing junctions by identifying cyclic node connections and we extracted the exact junction sequences absent from the previous publication, making them candidates for further biological validation (Supplemental Figs. S6–S7). We also overlaid miRNA site predictions (Dudekula et al. 2016). Figure 2A shows the E5–E2 back-splice junction with a predicted miRNA site in exon 4. We then examined the *HM13*::*TPX2* fusion from CCLE/DepMap (Bessière et al. 2024), where querying the breakpoint sequence retrieved the junction node in CCLE RNA-seq and confirmed expression specificity to the fusion-positive sample (Figure 2B). Long-read evidence further supported other breakpoints (Supplemental Fig. S5B). Vizitig graphs can also combine transcripts from several species, as illustrated for human *PAX6* and mouse *Pax6* (Supplemental Fig. S9), and provide a sequence-level alternative to classical splicing graphs (Supplemental Fig. S11).

**Figure 2:**
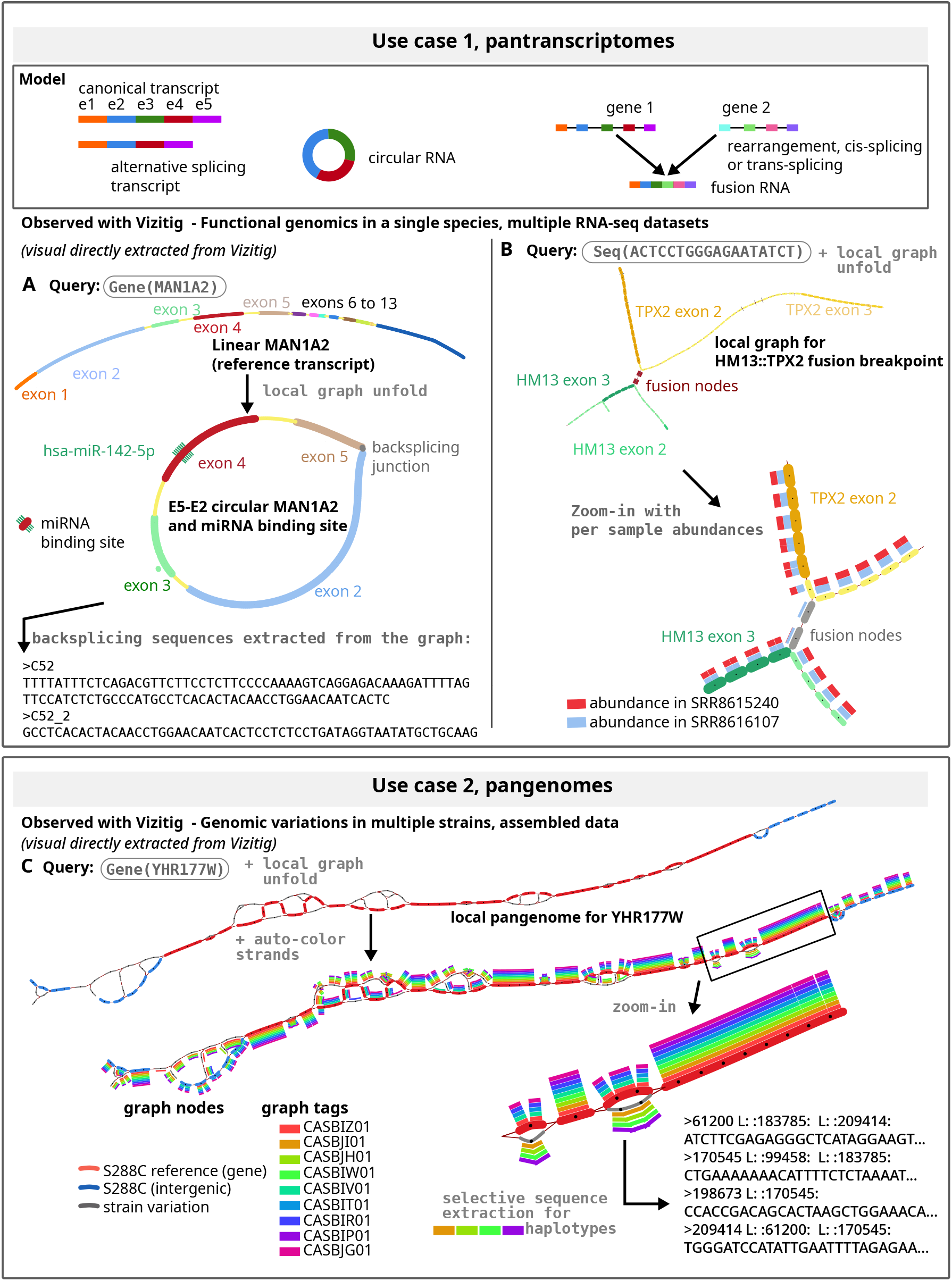
Illustrative examples of Vizitig use cases. **A** Circular RNA detection: visualization of linear and circular *MAN1A2* isoforms, including the E5–E2 back-splice junction and a predicted miRNA site on exon **B** Fusion exploration: a sequence query recovered the *HM13*::*TPX2* breakpoint, highlighting sample specificity. **C** Yeast pangenome: close-up of *S. cerevisiae YHR177W*. The top panel shows the reference (red), variants across nine strains (grey), and intergenic regions (blue). The bottom panel colors nodes by strain, revealing shared reference-like regions and specific variants. Variant sequences can be extracted directly for downstream analysis. All visualizations and colorings shown in A–C are exported directly from Vizitig as SVG.

### Exploration of pangenomes

We next turned to pangenomics, applying Vizitig across genomes of different size and repeat complexity: a human pangenome, a yeast pangenome, a rice pangenome and a *C. elegans* pangenome to illustrate scalability and core features.

We built a pangenome graph combining the *S. cerevisiae* reference strain S288C (∼12 Mb), nine telomere-to-telomere assemblies (O’Donnell et al. 2023), and GFF3 annotations, spanning ten strains in total. We queried the *YHR177W* gene-ID, reported to carry structural variations (O’Donnell et al. 2023), and showed complexity ranging from local synteny to inter-strain differences and conserved segments (Figure 2C, Supplemental Fig. S1). Vizitig can narrow the view to a selected subset of strains, and allows selective extraction of graph segments and haplotypes (Supplemental Material). We obtained similar results for *YDR018C* (Supplemental Fig. S4).

To demonstrate scalability to larger genomes, we constructed a pangenome of *Oryza sativa* ssp. *japonica* (rice, ∼380 Mb per assembly) using the Nipponbare reference genome (Kawahara et al. 2013) (IRGSP-1.0) and 10 chromosome-level cultivar assemblies from NCBI (accessions in Supplemental Table S2). The resulting de Bruijn graph (*k*=61) spans approximately 3.8 Gb of input sequence. We annotated the graph with Nipponbare gene models and exon sequences from the *Wx* (Waxy) gene, which controls amylose content in rice grains. A well-characterized G→ T single nucleotide polymorphism at the 5′ splice donor site of *Wx* intron 1 (Hirano et al. 1998) distinguishes the *Wx*^a^ allele (high amylose, non-sticky grain) from the *Wx*^b^ allele (low amylose, sticky grain), which predominates in temperate japonica cultivars. Using Vizitig, we queried the *Wx* locus and visualized the splice-site variant across all cultivars, confirming that sticky cultivars (Koshihikari, Hitomebore, Kitaake, Zhonghua 11) share the *Wx*^b^ allele, while tropical/ upland cultivars (Azucena, Kinandang Patong) carry the *Wx*^a^ allele. Two additional SNPs specific to the non-sticky phenotype were also identified in the graph (Supplemental Fig. S10). A larger *k* value (61 versus 31) was used for this genome to reduce spurious connections between paralogous gene family members, illustrating the importance of *kC*selection when working with repeat-rich genomes. We also illustrate a complementary use case with a *C. elegans* pangenome built from three chromosome-level assemblies (Bristol N2, Hawaiian CB4856, and DL226), revealing inter-strain structural variants on Chromosome I (Supplemental Fig. S8).

### Comparison with variation graph tools on human data

To further demonstrate the scalability of Vizitig, we compared it to variation graph tools on human data. Pangenome tooling remains fragmented: existing tools specialize in either graph construction, visualization, or querying, but none integrates all four functionalities. Figure 3 maps the current landscape of variation graph and de Bruijn graph tools across graph construction, annotation, querying, and visualization. While variation graph builders (Minigraph (Li et al. 2020), Minigraph-Cactus (Hickey et al. 2024), PGGB (Garrison et al. 2024)) and visualizers (Bandage (Wick et al. 2015) and BandageNG (https://github.com/asl/BandageNG), Sequence Tube Map (Beyer et al. 2019), VRPG (Miao and Yue 2025), odgi (Guarracino et al. 2022)) each address individual tasks, and de Bruijn graph builders (BCALM2 (Chikhi et al. 2016), GGCAT2 (Cracco and Tomescu 2023), Cuttlefish 3 (Khan et al. 2025)) focus on construction and ownership queries (i.e., whether a *k*-mer or a sequence appears in a color of the graph), Vizitig is the only tool that spans all four categories within a single pipeline, by integrating GGCAT.

**Figure 3:**
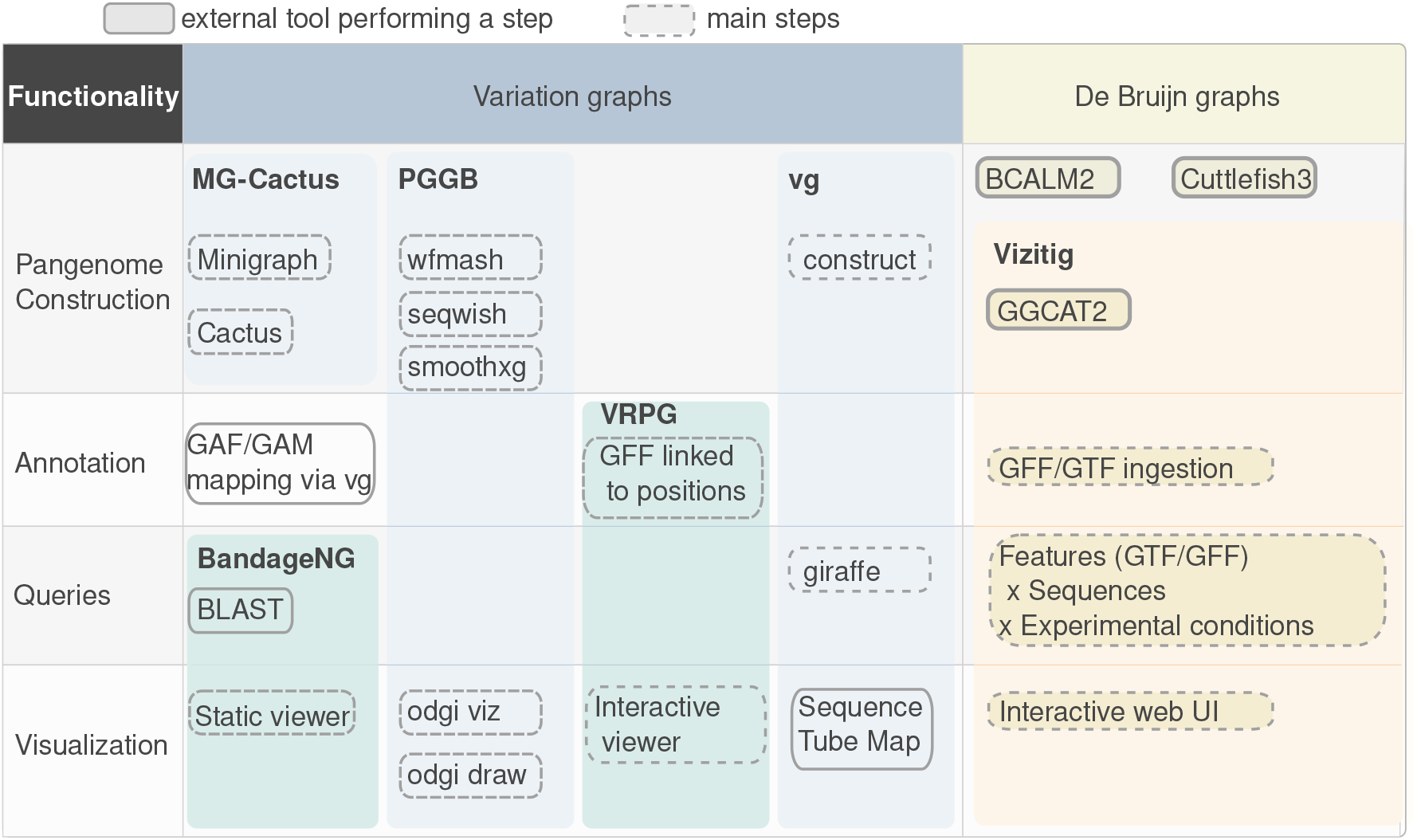
Pangenome graph tools landscape. Tools are positioned by graph type (variation graphs versus de Bruijn graphs) and functionality (construction, annotation, querying, visualization). Solid borders indicate external tools; dashed borders indicate internal features or subcommands. Nested bubbles indicate embedded components. Vizitig is the only tool that spans all four functional categories within a single pipeline.

To illustrate how these differences translate in practice, we compared how each builder and its associated visualization tool renders the same genomic region: a ∼3 kb window around exon 12 of the *DYRK1A* gene on Chr21, which contains several SNPs between the two HG002 haplotypes and the GRCh38 reference (Figure 4). Querying methods vary across tools: Vizitig allows direct access by gene or exon name, Bandage (Wick et al. 2015) requires BLAST (Altschul et al. 1990) in exact query mode, and Sequence Tube Map (Beyer et al. 2019) and odgi (Guarracino et al. 2022) require the exact genomic coordinates of the region.

**Figure 4:**
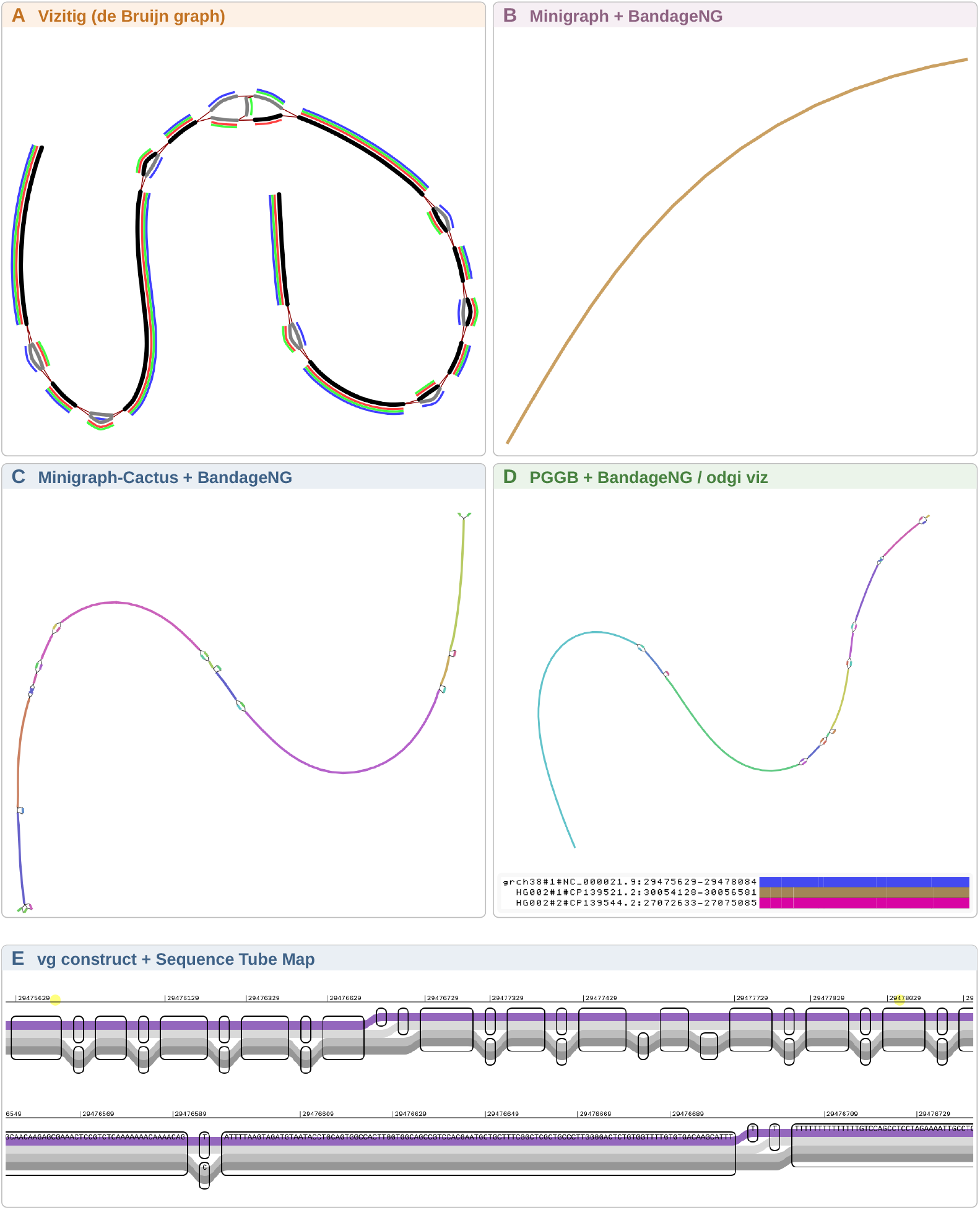
Comparison of visualization approaches. We worked on a ∼3 kb region around exon 12 of *DYRK1A* (Chr21) from the HG002 pangenome (2 haplotypes + GRCh38). **A** Vizitig: 2 commands + 1 query (`Gene(DYRK1A`)), haplotypes colored automatically. **B** Minigraph + BandageNG: captures SVs only (≥50 bp), no SNPs visible, requires BLAST queries. **C** Minigraph-Cactus + BandageNG: 3-step pipeline; base-level GFA must be identified and decompressed, requires BLAST queries. **D** PGGB + BandageNG / odgi viz: requires format conversion, coordinate lookup, region extraction, and sorting; 1D odgi visualization cannot resolve individual SNPs at this scale, 2D visualization could not be rendered on this example. **E** vg (Garrison et al. 2018) construct + Sequence Tube Map: requires a VCF file and multiple preprocessing steps; shows variant bubbles.

To quantify how Vizitig compares on human-scale data, we benchmarked per-chromo-some pangenome construction using two assembled haplotypes from the HG002 Ashkenazi trio (maternal and paternal) and the GRCh38 reference genome. Figure 5 shows wall time and peak RAM for Chr21 (∼46 Mb per haplotype) and Chr1 (∼248 Mb per haplotype) across Vizitig, Minigraph, Minigraph-Cactus, and PGGB. Direct performance comparisons should be interpreted with caution, as the underlying graph models differ fundamentally in construction strategy, resolution level, and intended use cases.

**Figure 5:**
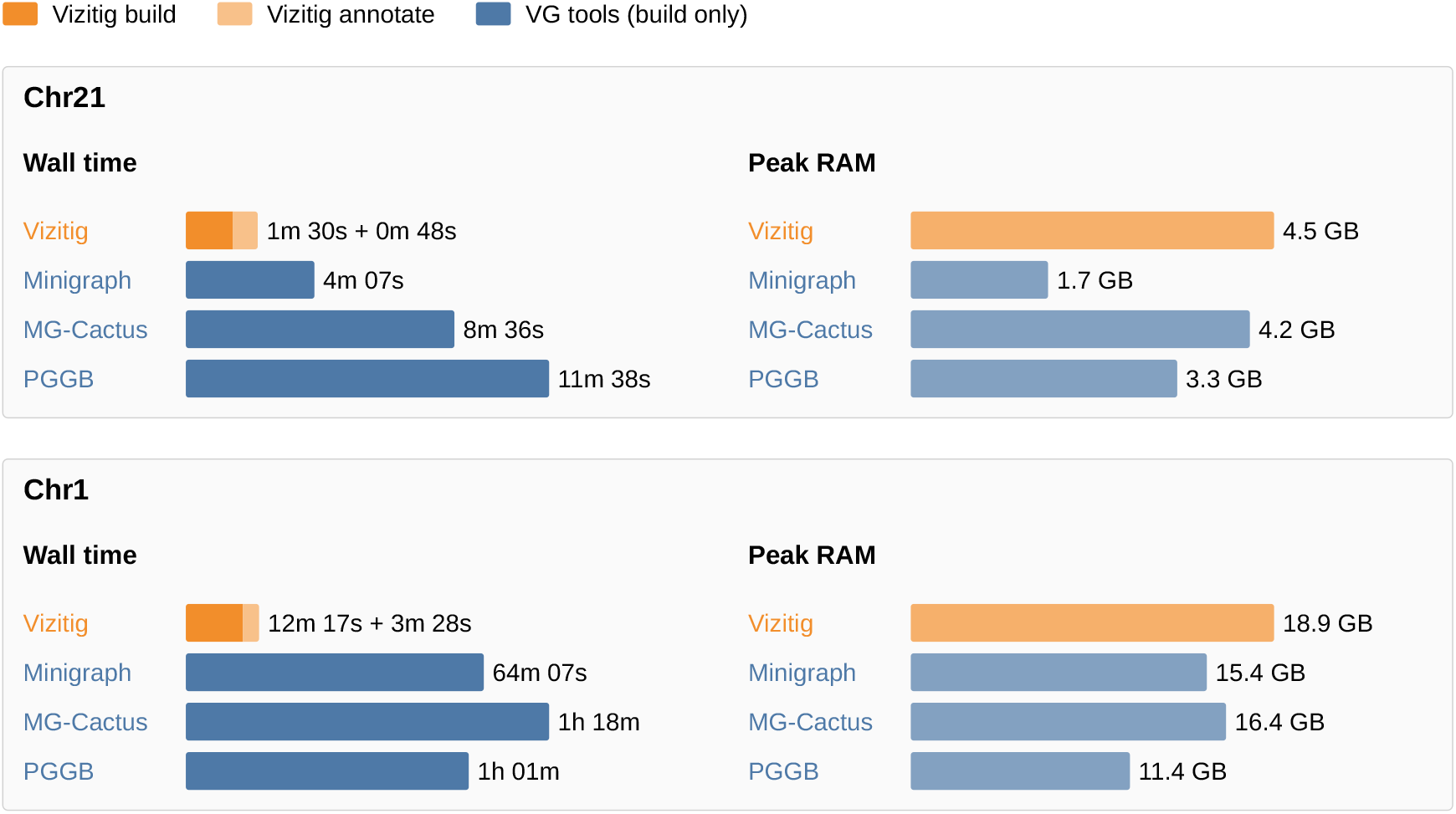
Per-chromosome human pangenome build resource comparison. On 2 HG002 haplotypes + GRCh38, Vizitig (*k*=61) time includes gene and exon annotation from GFF (MANE Select); stacked bars show build (dark) and annotation (light) time. Variation graph tools perform graph construction only. Minigraph-Cactus wall time is the sum of its three pipeline steps. Minigraph captures structural variants only (≥50 bp); Minigraph-Cactus and PGGB provide base-level resolution. Note that Vizitig’s peak RAM usage does not reflect a limitation, as the tool dynamically shards computation to stay within available memory (see Supplemental Table S5).

These applications highlight how Vizitig unifies queries, metadata, and visualization to support discovery and interpretation of complex sequence data, scaling from small microbial genomes to plant-scale and human pangenomes.

## Discussion

Vizitig provides the first unified platform for constructing, exploring, and querying de Bruijn graphs across multiple DNA and RNA samples. Through case studies in pantranscriptomics and pangenomics, we have demonstrated its ability to detect rare transcriptomic events such as fusion transcripts and circular RNAs, as well as to explore sequence variations and structural variations across yeast strains, nematode, rice cultivars and human pangenomes. The combination of an alignment-free index, feature-linked nodes, and on-demand subgraph loading distinguishes Vizitig from existing graph builders and viewers such as Bandage (Wick et al. 2015) and odgi (Guarracino et al. 2022), which primarily provide static visualization without integrated genomic feature, compositional querying or multi-sample visualization helpers.

A distinctive strength of Vizitig is its composable query language, which allows users to combine sequence motifs, genomic features, and experimental factors in a single expression. This goes beyond what any existing graph viewer or index offers: tools such as Bandage (Wick et al. 2015) require external BLAST queries and manual coloring to achieve what Vizitig expresses in one command, while large-scale *k*-mer indexes (Karasikov et al. 2025) support sequence membership queries but lack metadata composition and interactive visualization. The ability to formulate queries such as *retrieve a gene in a specific sample with abundance above a threshold* directly from the interface, without scripting or external tools, lowers the barrier for users who are not comfortable with complex command-line pipelines.

The alignment-free nature of de Bruijn graphs also brings practical advantages for pantranscriptomics. Because the graph is built from *k*-mer overlaps rather than from alignments to a reference, it captures all expressed sequences regardless of whether they appear in an annotation database. This is particularly valuable for detecting fusion transcripts and circular RNAs, which span non-canonical junctions absent from standard references. Our case studies illustrate that Vizitig can recover junction sequences that are absent from annotation data, such as fusions, opening the door to alignment-free detection of RNA variants.

From a practical standpoint, Vizitig is designed to run on commodity hardware: all experiments in this study, including the construction and exploration of human-scale pangenomes, were performed on a single laptop. The web-based interface requires no installation beyond the command-line tool itself, making it accessible to researchers without systems administration expertise. Pre-built databases can be shared as SQLite files, enabling collaborators to explore the same graphs without re-running the construction pipeline.

Our rice pangenome example (10 cultivars, ∼3.8 Gb total input) and our human pangenome examples (respectively 2 haplotypes + reference, ∼10 Gb and 10 samples, 20 haplotypes, ∼64 Gb total input) demonstrate that Vizitig scales to large-scale pangenomes. Our large-scale pantranscriptome (21 samples of human RNA-seq, 1,227 GB total input) demonstrates scalability on clinical-trial sized pantranscriptomes. A practical consideration for larger, repeat-rich genomes is the choice of *k*-mer size: larger values reduce spurious connections between paralogous sequences at the cost of increased graph fragmentation, while smaller values preserve more connectivity but may merge distinct loci. We found *k*=61 to be appropriate for plant genomes with moderate repeat content, and recommend users adjust this parameter based on genome complexity. Highly repetitive genomes (e.g., mammalian genomes) will require careful choice of *k*-mer size or chromosome-level partitioning to keep hub nodes interpretable, a trade-off each user will have to make according to the biological question at hand. For human data,

*k*=61 for genomic and *k*=31 for transcriptomic data are generally good values. Repeats are usually seen as a limitation of de Bruijn graphs, but their tendency to cluster together repeated sequences can also be an opportunity: for instance, Darmon et al. (2026) recently used de novo de Bruijn graph construction to annotate transposable elements directly from transcriptomic data, useful for non-model organisms lacking such resources. A current limitation is that Vizitig operates on de Bruijn graphs rather than accepting arbitrary pangenome graph formats. While this data-structure choice enables tight integration between graph construction, indexing, and querying, it means users with existing variation graphs must rebuild them as de Bruijn graphs. Another limitation is the absence of position-based queries: users cannot yet query by genomic coordinates on a reference, a feature that most genome browsers support and that would facilitate adoption by researchers accustomed to coordinate-based navigation. Supporting such queries requires mapping reference positions onto graph nodes, which is significantly more complex in de Bruijn graphs than in variation graphs where paths already carry coordinate information. Future work will focus on extending input format support, and enriching the query language to support position-based queries.

## Methods

### Software development and practices

Vizitig is an open-source software package designed for flexible deployment and integration in modern genomics workflows. It is distributed under the BSD license. The codebase is modular and actively maintained, with current test coverage exceeding 90%. The platform consists of a Python-based application programming interface (API), which enables advanced analytical operations such as graph-based filtering, locus comparison across samples, and metadata-driven search. This is complemented by a web-based frontend developed in JavaScript (using https://d3js.org/), designed for interactive visual exploration and intuitive manipulation of graph structures and sample metadata. The backend combines SQLite for portability and ease of deployment and Rust for performance-critical components such as *k*-mer indexing and query execution. This architecture ensures both speed and accessibility, making Vizitig usable across a range of computing environments, from personal workstations to cloud-based infrastructures. The frontend communicates via a lightweight web API with the core command-line engine, allowing researchers to query, visualize, and annotate graphs interactively. In addition, Vizitig supports export of graph views, metadata, and query results in standard formats such as GML, JSON, and SVG, export of subgraphs in BCALM2 format, and sharing of SQLite files themselves.

### Graph model

At the core of Vizitig is a graph-based representation of sequence data built on the de Bruijn graph model, a widely used structure in genome assembly and for alignment-free data representation. De Bruijn graphs capture the global structure of sequences, including chromosomes, transcripts, and variants, by representing *k*-mer overlaps between reads or assembled genomes. These graphs naturally encode biological variation through topological features such as branching paths and “bubbles” (Chikhi et al. 2016). Formally, given a set of sequences and a fixed integer *k*, the de Bruijn graph is defined as *G* = (*V, E*) where *V* is the set of all distinct *k*-mers (length-*k* substrings) occurring in the input sequences, and *E* ⊆ *V* × *V* is the set of edges: (*u, v*) belongs to *E* whenever the length-(*k* − 1) suffix of *u* equals the length-(*k* − 1) prefix of *v*, i.e., whenever *u* and *v* occur consecutively, overlapping on *k* − 1 nucleotides, in some input sequence. Since DNA is double-stranded, each *k*-mer *u* and its reverse complement *ū* represent the same underlying genomic locus; *k*-mers are therefore stored and indexed in canonical form, defined as min(*u, ū*) under a fixed ordering (e.g., lexicographic), so that a *k*-mer and its reverse complement are treated as a single node regardless of which strand a read originates from. In practice, Vizitig operates on the *compacted* de Bruijn graph, in which every maximal chain of *k*-mers with in-degree and out-degree equal to one is merged into a single node called a *unitig* (Chikhi et al. 2016). This lossless compaction drastically reduces the number of nodes and edges while preserving the full topology and sequence content of the graph.

To handle multi-sample datasets and support comparative analysis, we employ colored de Bruijn graphs (Marchet et al. 2021), an extension in which each node *v* ∈ *V* is additionally labeled with a color set *C*(*v*) ⊆ *S*, where *S* is the set of input samples (raw reads, assemblies, or reference sequences), indicating which samples contain that *k*-mer. This approach allows scalable co-integration of raw sequencing reads from different samples and reference sequences into a single graph, enabling more complete interpretation of partially covered or noisy regions.

Each unitig node stores an abundance value, computed as the average abundance of its constituent *k*-mers. These values provide an estimate of local sequence coverage and can be used directly in transcriptomic applications as a proxy for transcript-level expression (Bessière et al. 2024).

Integrating raw read datasets from diverse sources inevitably introduces noise, which can complicate analysis and interpretation. Vizitig addresses this challenge both upstream and downstream. During graph construction, an optional low frequency *k*-mer filtering step reduces sequencing artifacts and low-coverage errors, ensuring that the resulting graphs remain biologically meaningful and robust while accommodating heterogeneous input formats. Downstream, the visualization module includes an automated noise-filtering mechanism inspired by assembly pipelines (Bankevich et al. 2012), which prunes tips, i.e., short dead-end paths of low-abundance nodes typically arising from sequencing errors, along with other low-coverage artifacts unlikely to reflect true biological variation. A second feature can dynamically hide nodes or edges on demand, enabling users to focus on the relevant and biologically significant portions of the graph. Supplemental Fig. S12 shows how upstream and downstream filtering impacts a graph.

A known parameter sensitivity in de Bruijn graph construction is the choice of *k*-mer size, which can influence graph resolution and connectivity. In our experiments, we adopted standard *k*-mer sizes (from 21 to hundreds) that are commonly used in genome and transcriptome studies. To illustrate the robustness of Vizitig, we performed additional experiments (Supplemental Fig. S3) demonstrating that the biological signal of interest remains detectable even when different *k* values are applied.

### Database design

Vizitig’s backend currently utilizes SQLite (https://www.sqlite.org/) to manage unitig-defined graph structures, associating metadata directly with nodes and therefore allowing SQL queries on the graph. It incorporates sequences by marking nodes containing corresponding *k*-mers with relevant metadata from DNA/RNA sequences or annotations (GTF/GFF), and supports additional metadata such as color (sample identification) and abundance (occurrence in samples). Efficient retrieval is ensured through indexing, providing logarithmic query time for single *k*-mers and sequences of arbitrary lengths. Small-*k* indexes can be built to support fast querying of sequences smaller than *k*, but a linear fall-back exists that does not require re-building an index. A Python-based API and command-line interface offer comprehensive backend access for data manipulation. We chose not to rely on analytical graph databases such as Neo4j, as no mature, open-source, embedded solutions currently exist that can run on standard machines. Additionally, such databases are typically not optimized for rapid retrieval of localized graph regions, our primary query focus. To ensure optimal performance, we developed ad hoc indexes tailored for fast localized queries and integrated them with NetworkDisk (https://networkdisk.inria.fr/), a disk-based Python module compatible with NetworkX, facilitating large-scale graph manipulation with minimal memory usage by loading only relevant subsets during exploration.

### Human-based queries and interactions

A human-readable domain-specific query language (DSL), supported by a Python-based query assistant, translates user queries into executable NetworkDisk commands, facilitating graph manipulation. Additional DSL-driven filtering and transformation tools allow targeted data extraction and visualization. Vizitig’s domain-specific query language allows one to build queries that can be executed both as a retrieval query and as a filter query. The former fetches sequences and metadata from the database, while the latter applies visual transformations on data once it was retrieved. The domain-specific query language is composed of logical operators (AND, OR and NOT), parentheses, and tokens. Tokens are made of a type keyword, open parenthesis, an optional key and a closing parenthesis. Type keywords correspond to the different types of features and are generated based on the annotation files (e.g. `Gene, Transcript, Exon`). The specific keyword `Expe` allows a second, optional argument `A` that comes with an operator (`<, >, <=, >` = or =) and a value, that allows querying abundances. Querying by genomic coordinates is not yet supported (see Discussion).

Trait-based queries typically involve commands such as `Gene, Transcript`, or `Exon` to load regions of interest, often combined with abundance thresholds or sample presence. For example, `Expe(Sample_SRR8967009, A > 600`) outputs sequences from the given sample with abundance over 600. Sequence-based queries can be further refined with any available metadata or features, providing a flexible way to zoom into specific biological events.

Users can also alter the appearance of the graph to highlight specific samples, genes, or exons using custom colors and node shapes. We also implemented an autocolor tool that allows genes, exons, or chromosomes to be distinguished automatically, reducing the burden of repetitive manual operations. A built-in *sashimi-like* (Katz et al. 2015) visualization tool displays abundance levels at individual nodes across multiple samples within the graph (this tool produces results consistent with established quantification methods). The alignment module inside Vizitig implements the Smith-Waterman algorithm for sequence alignment.

## Performance

We collected the data availability, graph information, and performance for each experiment in Supplemental Tables S1–S2.

All Vizitig builds took less than 20 GB of RAM during construction for the different experiments. The query times for the different queries described in the article range from milliseconds to a maximum of 6 seconds of response time. We built de Bruijn graphs using BCALM2 (Chikhi et al. 2016) output format.

An extended version of the *HM13*::*TPX2* fusion experiment using 21 human RNA-seq datasets from SRA provides a view of Vizitig’s management of large workloads. All decompressed SRR reads totalled 1,227 gigabytes on disk. The references include hg38 RefSeq transcript sequences associated with hg38 reference metadata.

All experiments in this work were conducted on a laptop (CPU Intel Core Ultra 7 165H processor with 16 cores and 22 logical threads at 5.0 GHz, RAM 32 GB of DDR5-5600 (2×16 GB modules) at 5600 MT/s, Disk 4.0 TB NVMe SSD with 512-byte logical block addressing, mounted as the secondary data storage partition). The graph was built using BCALM2 (*k*=31, minimum abundance = 2, final size on disk 67.5 GB, index size on disk 13.8 GB).

Despite data size largely exceeding RAM capacity, Vizitig is capable of managing the workload by chunking data and sharding the graph using its indexes. The coloring phase time is linear in number of samples. Peaks in CPU usage are caused by index join operations. Peaks in RAM usage usually follow, depending on the size of the join result. In both cases, the workload is distributed to prevent resource exhaustion that would lead to system interruption (see complete resource monitoring in Supplemental Table S4).

### Use cases

#### Pantranscriptomes and rare variants

Circular RNAs *MAN1A2* and *FBXW4* were previously described by Mercer et al. (2015). In that study, circRNAs were identified through total RNA extraction, poly(A) depletion to enrich for pre-mRNA (including introns and lariats), RNase R digestion of linear RNA, and subsequent validation by sequence mapping. Using Vizitig, we located alternative circular back-splicing junctions for both genes by querying nodes connecting the reported junctions and extracted the corresponding sequences, which had not been documented in the original publication. To demonstrate integration of external data, we incorporated predictions from circInteractome (Dudekula et al. 2016), which identifies miRNA binding sites on circRNAs. For *MAN1A2*, predicted sites were matched to the graph through sequence queries, and conserved sites were located within the Vizitig graph. Additional *MAN1A2* and *FBXW4* junctions appear in Supplemental Figs. S6 and S7.

We next investigated the *HM13*::*TPX2* human gene fusion transcript, reported in the Cancer Cell Line Encyclopedia (CCLE) and DepMap/COSMIC fusion database (Bessière et al. 2024).

Fusion junction coordinates were obtained from STAR-Fusion (Gillani et al. 2021) alignments provided by DepMap, following guidelines to select events with high reliability. A de Bruijn graph (*k*=61) was constructed from two control RNA-seq datasets (SRR8615240, SRR8615242) and one dataset known to harbor the fusion event (SRR8616107). Reference transcripts and abundance data were annotated similarly to the previous example. Using the fusion breakpoint sequence as a locus query within Vizitig, capable of querying sequences both shorter and longer than the *k*-mer size, we identified the exact node corresponding to the fusion junction (Figure 2B). This node, not associated with known genes or annotations, remained distinctly uncolored. Applying the `Sashimi` action revealed expression specificity of the fusion to the relevant dataset (Figure 2B), confirming its sample-specific presence. Beyond single events, Vizitig allows interactive expansion of the graph around a locus to reveal neighboring structures, exemplified by the detection of an additional *BCL2L1*::*HM13* fusion in the same dataset, also reported in DepMap (Supplemental Fig. S5A). Building on this, we examined the *CYTH1*::*EIF3H* event in the SKBR3 cell line, previously characterized by Qin et al. (2025). Although that study combined long- and short-read sequencing, the fusion sequences themselves were not directly available. Vizitig enabled their retrieval and visualization, showing long-read support for the breakpoint and identifying bridging nodes between the two genes, including one of the reported breakpoints (Supplemental Fig. S5B).

### Extraction of sequences of interest in pangenomics

We constructed a yeast pangenome by combining the *S. cerevisiae* reference strain S288C (with annotations) and nine additional assemblies from a recent study of 142 high-quality, telomere-to-telomere genomes (O’Donnell et al. 2023), which characterized structural variation among strains. Annotation data from the S288C GFF3 file were incorporated to enable direct feature queries. We selected the *YHR177W* gene, reported to carry multiple structural variants in the original study, and demonstrated the levels of complexity Vizitig can expose, from a linear view of the gene to inter-strain variation and conserved regions. Vizitig can automatically display strain support for variants (critical when analyzing many strains) or restrict the view to a subset of interest, focusing on bio-logically meaningful differences. It also supports selective extraction of graph segments and sequences. As an example, we selected a strain with documented structural variants, visually confirmed in Vizitig, and extracted the sequences corresponding to its haplotype (see Supplemental Material). The *YDR018C* example (Supplemental Fig. S4) and the *C. elegans* Chromosome I pangenome (Supplemental Fig. S8) were built following the same procedure; assembly accessions are given in Supplemental Material.

All Vizitig graphs, Supplemental Material, and instructions to reproduce experiments are available at https://zenodo.org/records/22141527.

## Data sources

All data analyzed in this study were previously published and are publicly available; accession numbers for every experiment are listed in Supplemental Tables S1 and S2. RNA-seq reads were downloaded from the NCBI Sequence Read Archive (SRA; https://www.ncbi.nlm.nih.gov/sra). Genome assemblies (rice cultivars, *C. elegans* strains, and the GRCh38 reference) were retrieved from NCBI Datasets (https://www.ncbi.nlm.nih.gov/datasets/genome/); *S. cerevisiae* telomere-to-telomere assemblies from the European Nucleotide Archive (ENA; https://www.ebi.ac.uk/ena/browser/, project PRJEB59413), and the reference strains used in Supplemental Fig. S2 from the Saccharomyces Genome Database (SGD; https://www.yeastgenome.org/). HG002 assemblies were obtained from the Human Pangenome Reference Consortium (HPRC; https://humanpangenome.org/) and the Genome in a Bottle Consortium (GIAB; https://www.nist.gov/programs-projects/genome-bottle). Human transcript sequences and annotations were obtained from NCBI RefSeq (https://www.ncbi.nlm.nih.gov/refseq/), including MANE Select (https://www.ncbi.nlm.nih.gov/refseq/MANE/), fusion junctions from DepMap (https://depmap.org/portal/), and the hsa-miR-142-5p sequence from miRBase (https://www.mirbase.org/).

## Software availability

Vizitig is available as open-source software under the BSD license. The source code, documentation, and issue tracker can be found at: https://gitlab.inria.fr/vizisoft/vizitig. A frozen snapshot of the source code and example scripts used in this study is provided as Supplemental Code.

## Acknowledgments

This work is funded by ANR Find-RNA ANR-23-CE45-0003.

The authors would like to thank Rémi Morvan, Margaux Mouton, Loup Lobet, Cheyma Karbiche, Loïc Marmey and Gloria Kristal Medina Vargas who helped test Vizitig v0.* to 1.0, and Bruno Guillon for his help regarding NetworkDisk. The authors address their warmest regards to Jia-Xing Yue and Miao Zepu for allowing to use their figure from the VRPG paper. They would like to thank proofreaders including Thérèse Commes, Rayan Chikhi, and Lucien Piat who helped greatly with the manuscript.

## Author Contributions

B.D. developed the software and performed experiments. C.P. contributed to algorithm design. C.M. conceived and supervised the project. All authors wrote the manuscript.

## References

Altschul SF, Gish W, Miller W, Myers EW, Lipman DJ. 1990. Basic local alignment search tool. J Mol Biol 215: 403–410.

Bankevich A, Nurk S, Antipov D, Gurevich AA, Dvorkin M, Kulikov AS, Lesin VM, Nikolenko SI, Pham S, Prjibelski AD, et al. 2012. SPAdes: a new genome assembly algorithm and its applications to single-cell sequencing. J Comput Biol 19: 455–477.

Bessière C, Xue H, Guibert B, Boureux A, Rufflé F, Viot J, Chikhi R, Salson M, Marchet C, Commes T, et al. 2024. Transipedia.org: k-mer-based exploration of large RNA sequencing datasets and application to cancer data. Genome Biol 25: 266.

Beyer W, Novak AM, Hickey G, Chan J, Tan V, Paten B, Zerbino DR. 2019. Sequence tube maps: making graph genomes intuitive to commuters. Bioinformatics 35: 5318–5320.

Chikhi R, Dufresne Y, Medvedev P. 2024. Constructing and personalizing population pangenome graphs. Nat Methods 21: 1980–1981.

Chikhi R, Limasset A, Medvedev P. 2016. Compacting de Bruijn graphs from sequencing data quickly and in low memory. Bioinformatics 32: i201–i208.

Cracco A, Tomescu AI. 2023. Extremely fast construction and querying of compacted and colored de Bruijn graphs with GGCAT. Genome Res 33: 1198–1207.

Darmon S, Mary A, Lacroix V. 2026. Extended t-cores for the de novo identification of transposable elements and other inexact repeats from short read RNA-seq data. bioRxiv doi: 10.64898/2026.07.06.736737.

Dudekula DB, Panda AC, Grammatikakis I, De S, Abdelmohsen K, Gorospe M. 2016. CircInteractome: a web tool for exploring circular RNAs and their interacting proteins and microRNAs. RNA Biol 13: 34–42.

Gao Y, Yang X, Chen H, Tan X, Yang Z, Deng L, Wang B, Kong S, Li S, Cui Y, et al. 2023.A pangenome reference of 36 Chinese populations. Nature 619: 112–121.

Garrison E, Guarracino A, Heumos S, Villani F, Bao Z, Tattini L, Hagmann J, Vorbrugg S, Marco-Sola S, Kubica C, et al. 2024. Building pangenome graphs. Nat Methods 21: 2008–2012.

Garrison E, Sirén J, Novak AM, Hickey G, Eizenga JM, Dawson ET, Jones W, Garg S, Markello C, Lin MF, et al. 2018. Variation graph toolkit improves read mapping by representing genetic variation in the reference. Nat Biotechnol 36: 875–879.

Gillani R, Seong BKA, Crowdis J, Conway JR, Dharia NV, Alimohamed S, Haas BJ, Han K, Park J, Dietlein F, et al. 2021. Gene fusions create partner and collateral dependencies essential to cancer cell survival. Cancer Res 81: 3971–3984.

Guarracino A, Heumos S, Nahnsen S, Prins P, Garrison E. 2022. ODGI: understanding pangenome graphs. Bioinformatics 38: 3319–3326.

Hickey G, Monlong J, Ebler J, Novak AM, Eizenga JM, Gao Y, Human Pangenome Reference Consortium, Abel HJ, Antonacci-Fulton LL, Asri M, et al. 2024. Pangenome graph construction from genome alignments with Minigraph-Cactus. Nat Biotechnol 42: 663–673.

Hirano H-Y, Eiguchi M, Sano Y. 1998. A single base change altered the regulation of the Waxy gene at the posttranscriptional level during the domestication of rice. Mol Biol Evol 15: 978–987.

Holley G, Eggertsson HP, Kristmundsdottir S, Beyter D, Skuladottir AT, Moore KH, Olason PI, Gylfason A, Magnusson OT, Oddsson A, et al. 2026. An Icelandic pangenome reference. Nature doi: 10.1038/s41586-026-10924-7.

Jonkheer EM, van Workum D-JM, Sheikhizadeh Anari S, Brankovics B, de Haan JR, Berke L, van der Lee TA, de Ridder D, Smit S. 2022. PanTools v3: functional annotation, classification and phylogenomics. Bioinformatics 38: 4403–4405.

Karasikov M, Mustafa H, Danciu D, Kulkov O, Zimmermann M, Barber C, Rätsch G, Kahles A. 2025. Efficient and accurate search in petabase-scale sequence repositories. Nature 647: 1036–1044.

Katz Y, Wang ET, Silterra J, Schwartz S, Wong B, Thorvaldsdóttir H, Robinson JT, Mesirov JP, Airoldi EM, Burge CB. 2015. Quantitative visualization of alternative exon expression from RNA-seq data. Bioinformatics 31: 2400–2402.

Kawahara Y, de la Bastide M, Hamilton JP, Kanamori H, McCombie WR, Ouyang S, Schwartz DC, Tanaka T, Wu J, Zhou S, et al. 2013. Improvement of the Oryza sativa Nipponbare reference genome using next generation sequence and optical map data. Rice 6: 4.

Khan J, Dhulipala L, Pandey P, Patro R. 2025. Faster and Scalable Parallel External-Memory Construction of Colored Compacted de Bruijn Graphs with Cuttlefish 3. bioRxiv doi: 10.1101/2025.02.02.636161.

Leonard AS, Mapel XM, Pausch H. 2024. Pangenome-genotyped structural variation improves molecular phenotype mapping in cattle. Genome Res 34: 300–309.

Li H, Feng X, Chu C. 2020. The design and construction of reference pangenome graphs with minigraph. Genome Biol 21: 265.

Liao W-W, Asri M, Ebler J, Doerr D, Haukness M, Hickey G, Lu S, Lucas JK, Monlong J, Abel HJ, et al. 2023. A draft human pangenome reference. Nature 617: 312–324.

Marchet C, Boucher C, Puglisi SJ, Medvedev P, Salson M, Chikhi R. 2021. Data structures based on k-mers for querying large collections of sequencing data sets. Genome Res 31: 1–12.

Mercer TR, Clark MB, Andersen SB, Brunck ME, Haerty W, Crawford J, Taft RJ, Nielsen LK, Dinger ME, Mattick JS. 2015. Genome-wide discovery of human splicing branch-points. Genome Res 25: 290–303.

Miao Z, Yue J-X. 2025. Interactive visualization and interpretation of pangenome graphs by linear-reference-based coordinate projection and annotation integration. Genome Res 35: 296–310.

O’Donnell S, Yue J-X, Saada OA, Agier N, Caradec C, Cokelaer T, De Chiara M, Delmas S, Dutreux F, Fournier T, et al. 2023. Telomere-to-telomere assemblies of 142 strains characterize the genome structural landscape in Saccharomyces cerevisiae. Nat Genet 55: 1390–1399.

Qin Q, Popic V, Wienand K, Yu H, White E, Khorgade A, Shin A, Georgescu C, Campbell CD, Dondi A, et al. 2025. Accurate fusion transcript identification from long- and short-read isoform sequencing at bulk or single-cell resolution. Genome Res 35: 967–986.

Thorvaldsdóttir H, Robinson JT, Mesirov JP. 2013. Integrative Genomics Viewer (IGV): high-performance genomics data visualization and exploration. Brief Bioinform 14: 178–192.

Wick RR, Schultz MB, Zobel J, Holt KE. 2015. Bandage: interactive visualization of de novo genome assemblies. Bioinformatics 31: 3350–3352.

