## Supplementary material for "The Vizitig pangenome and pantranscriptome explorer": suppl_text_and_figures

#### Data and graphs

Details on data availability, graph characteristics, and performance for each experiment are provided in Supplemental Tables S1 and S2.

| **Study** | **Sequence accessions** | **Reference** | **Graph specs** | **Time for build** |
|---|---|---|---|---|
| *HM13*::*TPX2* extended (21 samples) | 10 fusion samples: SRR8615719, SRR8615900, SRR8615457, SRR8616176, SRR8615604, SRR8615901, SRR8615899, SRR8615456, SRR8616107, SRR8615602; 11 non-fusion samples: SRR8615240–SRR8615250 | hg38 RefSeq transcripts | $k$=31, 1,227 GB input, 67.5 GB graph, 13.8 GB index | 9h 23min (see Table S4) |
| *HM13*::*TPX2* fusion | SRR8616107, SRR8615240, SRR8615242 | hg38 RefSeq transcripts | $k$=61, 9.6M nodes, 5.6 GB | 15 min |
| *MAN1A2* and *FBXW4* circular RNAs | SRR1049830, SRR1049831, SRR1049832, SRR1049833 | hg38 RefSeq transcripts, hsa-miR-142-5p from miRBase | $k$=31, 30.7M nodes, 13.9 GB | 40 min |
| *CYTH1*::*EIF3H* fusion | SRR28000123_1, SRR28000123_2 (Illumina short reads), SRR28000125 (PacBio reads) | hg38 RefSeq transcripts | $k$=31, 15.2M nodes, 7.8 GB | 20 min |
| *PAX6*/*Pax6* pantranscriptome | – | NM_000280 and NM_001244198 from NCBI | $k$=31, 64 nodes, < 1 GB | < 1 min |

Table S1: Summary of datasets, references, graph specifications, and build times for Vizitig experiments (transcriptomics).

| Study | Sequence accessions | Reference | Graph specs | Time for build |
|---|---|---|---|---|
| *Saccharomyces cerevisiae* pangenome | CASBIP01, CASBIR01, CASBIT01, CASBIV01, CASBIW01, CASBJG01, CASBJH01, CASBJI01, CASBIZ01 from PRJEB59413 | S288C | $k$=41, 500k nodes, 136 MB | 5 min |
| *Oryza sativa* ssp. *japonica* pangenome (rice) | GCA_026167685.1 (Changxianggeng 1813), GCA_055690965.1 (Kinandang Patong), GCA_009830595.1 (Azucena), GCA_014526345.1 (Zhonghua 11), GCA_009797565.1 (Kitaake), GCA_000817635.1 (A123), GCA_000817615.1 (HEG4), GCA_000321445.1 (Hitomebore), GCA_000164945.1 (Koshihikari), GCA_055397395.1 (Udai21) | Nipponbare IRGSP-1.0 GCF_001433935.1 + gene annotation | $k$=61, $f$=1, ~3.8 Gb input | 14 min |
| Human HG002 trio (per-chromosome) | GCA_018852605.1, GCA_018852615.1 (GIAB), GRCh38 | GRCh38 + RefSeqGene GFF (MANE Select) | $k$=61, per-chromosome graphs | Chr21: 2m18s; Chr1: 15m45s (build+annot.) |
| Human pangenome – 1 sample (whole-genome) | HG002: GCA_018852605.3 (pat), GCA_018852615.3 (mat) from HPRC | – | $k$=61, 18.7M nodes, 2.9 billion $k$-mers, 16.9 GB graph, 87.5 GB index | 7h 34min (see Table S5) |
| Human pangenome – 10 samples (whole-genome) | HG002, HG005, HG00438, HG00621, HG00673, HG00733, HG00735, HG00741, HG01071, HG01106 (20 haplotypes from HPRC) | – | $k$=61 | – (see Table S5) |

Table S2: Summary of datasets, references, graph specifications, and build times for Vizitig experiments (pangenomics).

| Tool | Version | Purpose |
|---|---|---|
| Vizitig | 1.0.5 | Pangenome/pantranscriptome construction, indexing, and visualization |
| GGCAT | 2.0.0 | Compacted de Bruijn graph construction (embedded in Vizitig) |
| BCALM2 | 2.2.3 | Compacted de Bruijn graph construction (legacy) |
| Minigraph | 0.21-r606 | Variation graph construction |
| Minigraph-Cactus | 9.4.1 | Variation graph construction |
| PGGB | 0.7.4 | Variation graph construction |
| BandageNG | 2026.6.1 | Graph visualization (comparison) |
| odgi | 0.9.4 | Graph toolkit and visualization (comparison) |
| Sequence Tube Map | vg1.48.0 (Docker) | Graph visualization (comparison) |
| vg | 1.75.1 | Variation graph toolkit |

Table S3: Software versions used in this study.

#### Performance

Table S4 provides resource usage plots for running Vizitig in the largest experiment in this work, an extension of the *HM13*::*TPX2* dataset (first row of Table S1), using 21 human RNA-seq SRA datasets. The graph was built using BCALM2 ($k$=31, minimum abundance = 2, final size on disk 67.5 GB, index size on disk 13.8 GB).

| | Time | CPU usage | RAM usage |
|---|---|---|---|
| Build and index | 1h15min | 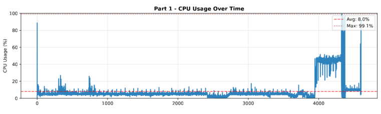 | 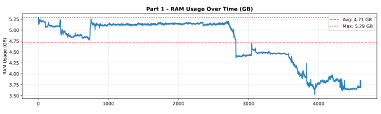 |
| Coloring graph | 7h20min | 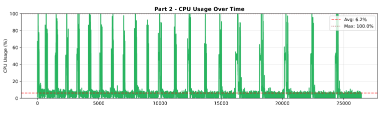 | 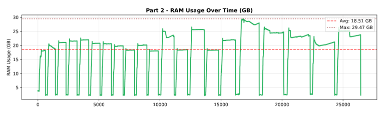 |
| Annotate | 48min | 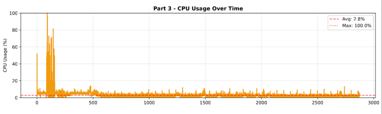 | 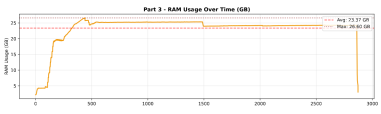 |

Table S4: Detailed performance of Vizitig on a large dataset. All measurements include the residual usage of CPU and RAM by the operating system. The build and index phase refers to the ingestion of data in the database and the indexing of $k$-mers. The "coloring graph" phase refers to tagging the graph with the origin sample of each $k$-mer and its abundance in its sample. The "annotate" phase refers to tagging $k$-mers if they are found in the reference sequences of the transcripts.

The analysis of performance and resource usage shows that Vizitig manages memory to handle large-scale tasks. The workload is balanced between available threads and sharded according to the available memory in order to prevent system crashes and allow scalability to large experiments. The CPU usage is not always high due to writing phases in the database: Vizitig's bottleneck is writing speed, since computation phases are highly optimized to leverage $k$-mer lookups, which are fast in our model.

#### Scalability on human pangenome data

To evaluate Vizitig's scalability on whole-genome human pangenome data, we benchmarked the full `vizitig create` pipeline (graph construction, indexing, and coloring) on assembled haplotypes from the HPRC HG002 Ashkenazi trio. All experiments were run on the same laptop as all other experiments (Intel Core Ultra 7 165H, 22 threads, 32 GB RAM, NVMe SSD storage). Table S5 reports wall time and resource usage for increasing numbers of haplotypes. The build time and indexing show sublinearity, meaning that they grow slower than the number of samples. This is expected in a de Bruijn graph, where the number of new $k$-mers added by a new sample is small when genomes are similar. The coloring phase shows superlinearity, which is also expected: as the number of nodes in the graph increases, the number of operations required to tag each node with its sample of origin grows accordingly. Additionally, when available memory is limited, the intersection between the $k$-mers of the current sample and the $k$-mers of the graph must be heavily sharded, causing the tagging operation to be repeated many times. This effect does not arise for smaller genomes such as bacterial pangenomes or single-chromosome human pangenomes. Furthermore, the current coloring implementation processes samples sequentially, and parallelization across samples would reduce the overall time. Alternatively, the coloring step could be replaced by exact matching of unitig sequences against the sample files, a highly efficient operation using standard Unix text-search utilities such as `ripgrep`, requiring no specialized bioinformatics software. In the case of large human pangenomes, the trade-off between precomputing and storing the origin of each $k$-mer versus searching for it on the fly in the sample files remains an open question. Overall, Vizitig scales linearly with the number of samples, whereas all-vs-all variation graphs scale quadratically.

| **Dataset** | **Haplotypes** | **Input size** | **Graph construction** | **Indexing** | **Coloring** | **Total time** |
|---|---|---|---|---|---|---|
| HG002 (1 sample) | 2 | 5.7 GB | 22 min | 42 min | 6h 29min | 7h 34min |
| HPRC (10 samples) | 20 | 57 GB | 1h 50min (5×) | 1h 11min (1.7×) | 150h (23×) | 153h (20×) |

Table S5: Scalability of `vizitig create` ($k$=61, $f$=1) on whole-genome human pangenome data. The 1-sample dataset consists of two assembled haplotypes (maternal and paternal) from the HPRC HG002 Ashkenazi trio. The 10-sample dataset uses 20 haplotypes from HPRC. Graph construction includes subgraph assembly (GGCAT), global graph merging, and database ingestion. Indexing builds the $k$-mer index (RustIndex). Coloring tags each $k$-mer with its sample of origin and abundance. Values in parentheses report the ratio relative to the 1-sample baseline.

### Supplemental results

#### Comparison to other graph visualizations

| Category | Capability | Bandage | Vizitig |
|---|---|---|---|
| Graph | Input | GFA | FASTA |
| | Output | – | export and selective export in BCALM, JSON, SVG |
| | Load subgraph | ✓ | ✓ |
| | Iterative exploration | – | ✓ |
| | Iterative exploration constrained by metadata | – | ✓ |
| | Orientation (bi-directional) | ✓ | ✓ |
| Queries | Query with a sequence | – | ✓ |
| | Query with metadata | – | ✓ |
| | Query with node ID | ✓ | ✓ |
| | Combination (AND, OR, …) of queries | – | ✓ |
| | Alignment | Built-in BLAST (to external database) | Built-in SW (for graph nodes) |
| Visualization | Zoom, rotate | ✓ | ✓ |
| | Node length as a function of sequence length | ✓ | ✓ |
| | Change graph layout (linear, circular) | – | ✓ |
| | User-defined colors and thickness | ✓ | ✓ |
| | Label nodes with annotation | Restricted to BLAST matches | Manual and automated, on any feature (gene, exon, chromosome, …) |
| | Thickness as a function of abundance (Sashimi-like) | ✓ | ✓ |

Table S6: Capabilities overview between Bandage and Vizitig.

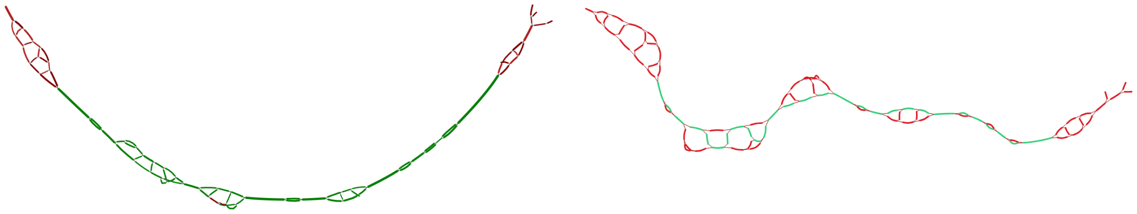

Figure S1: Comparison between the Bandage visualization (left) and Vizitig visualization (right) of the *YHR177W* gene in the pangenome of 10 *S. cerevisiae* assembled genomes. In green are the matches of the gene in BLAST for Bandage and the reference annotations of S288C for Vizitig.

List of the strains used to build this graph in Figure S1: CASBIH01, CASBIF01, CASBII01, CASBIP01, CASBIR01, CASBIS01, CASBIT01, CASBIV01, CASBIY01, and CASBJB01. The graph was built using BCALM2 and converted to GFA using the integrated script. The following actions are required to recreate the figures:

- **Bandage:** Load the graph, create a BLAST hit database using the sequence of the gene, query the graph using BLAST's matches at a distance of 6 nodes, set up color mode to BLAST's matches.
- **Vizitig:** Build the graph, annotate the graph with reference sequence and strains, query the gene (`Gene(YHR177W)`), color the gene in black, autocolor the strains by tagging the nodes.

The same patterns are shown in both visualizations. The gene of interest is colored green (left) and black (right) and three different complex components are displayed, as well as five bubble patterns depicting simple variation. Two complex components are found at the end of the matching nodes on both sides. In Bandage, the green nodes represent the zones of the graph that match with at least one BLAST reference. It is possible to configure the software to display the depth of the match. In Vizitig, the black nodes are found in the reference sequence, and each line of color represents a match of the sequence with one strain, making the study of inter-strain variations more natural.

Both Bandage and Vizitig can display similar graph patterns, such as the complex components and bubble variations observed around the *YHR177W* gene. However, Vizitig goes far beyond visualization: it acts as a query and exploration platform, whereas Bandage remains primarily a viewer. In Bandage, visualizing a gene requires several manual steps (building a BLAST database, querying the graph, and coloring the resulting matches) while Vizitig performs equivalent operations through integrated commands and metadata-aware queries.

Vizitig accepts a broad range of query types, including and combining node or gene identifiers, sequences, and metadata such as sample or condition, enabling users to explore multi-sample graphs in a single interface. Vizitig specializes in treating multi-sample input notably through automatic coloring and dynamic updates to highlight sample-specific variations. The visualizations also differ in dynamicity: Bandage regenerates an image after each new query, whereas Vizitig dynamically adds nodes to the loaded subgraph without regenerating static images. Vizitig's platform also provides direct access to node sequences and all associated metadata in an adjacent interactive table, while Bandage displays the node ID. Finally, in Vizitig, the layout of the nodes can be changed using several algorithms and the whole graph can be reconfigured manually.

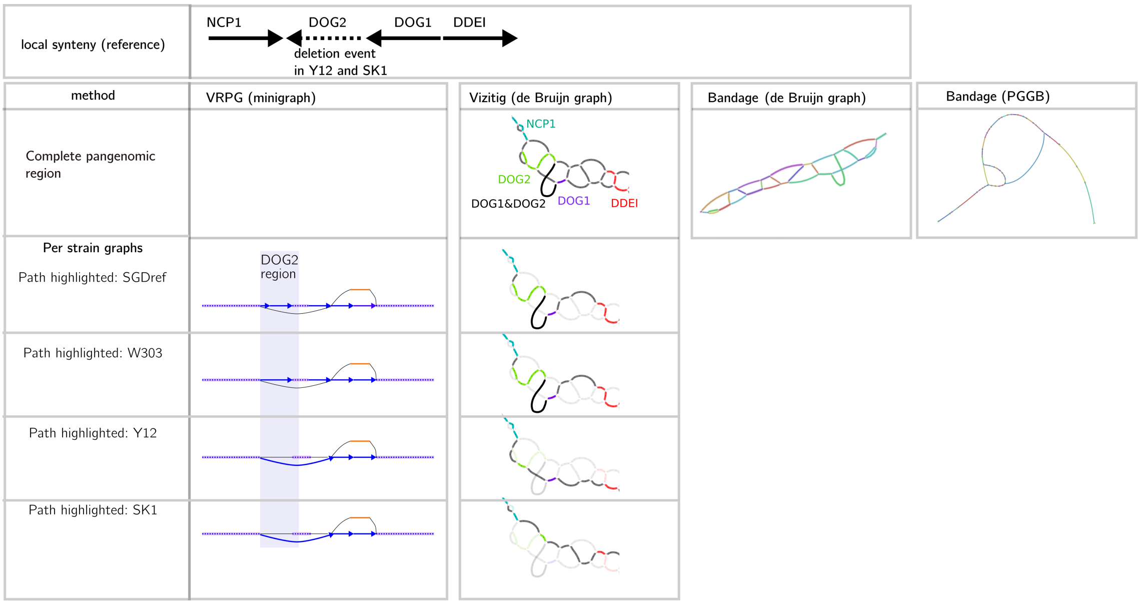


Figure S2: Graph model (variation graph and de Bruijn graph) and visualization (VRPG (Miao and Yue 2025), Vizitig, and Bandage (Wick et al. 2015)) comparison on four strains of *S. cerevisiae* on the well-documented deletion of the *DOG2* gene in the Y12 and SK1 strains. VRPG and Vizitig allow the individual highlight of strains. VRPG panel adapted from Miao and Yue (2025) (CC BY-NC 4.0) with permission from the authors.

We compare the de Bruijn graph and the variation graph (built using Minigraph (Li et al. 2020) and PGGB (Garrison et al. 2024)) of four different strains of *S. cerevisiae*. The genomes of the four reference strains were obtained from the Saccharomyces Genome Database (SGD; https://www.yeastgenome.org/). Unfortunately, the state of the VRPG release did not allow its proper functioning due to compilation and execution errors; therefore, the figure in the VRPG (minigraph) column is a manual reproduction of Figure 4 in the VRPG publication (Miao and Yue 2025).

In VRPG and Vizitig we highlighted paths for each strain, compared to the reference strain. The general variations between strains are shown in both visualizations, but the level of detail is higher in de Bruijn graphs, depicting intra-gene variations that cannot be seen in variation graphs (see genes *DOG1* and *DDEI* in Y12 or *NCP1* and *DOG1* in SK1 for instance). The visual transformations applied to the graph with Vizitig depict strain variations along with the genetic affiliation of the sequences.

In Bandage, it would be possible to produce the same strain variation result by creating a BLAST query with the sequence of each strain, expanding the neighborhood of the graph, and coloring the strain using BLAST hit coloring.

#### Graph model

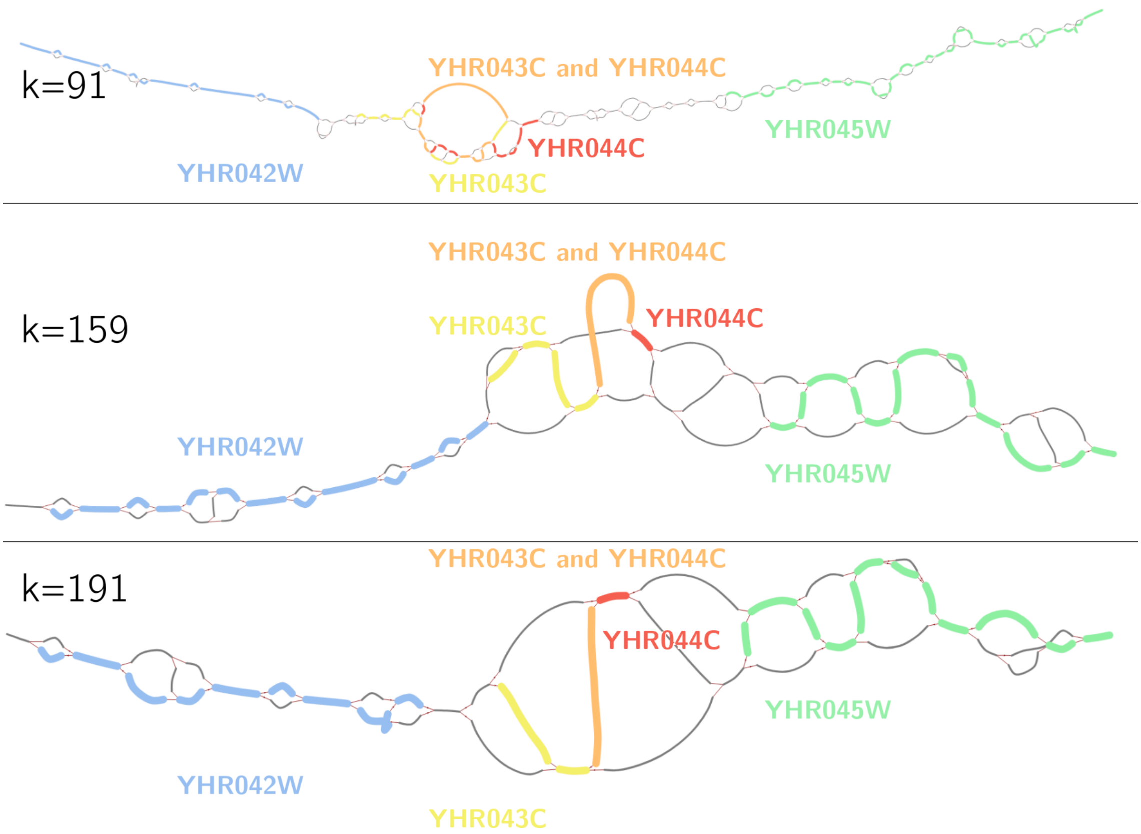


Figure S3: Pangenomes of four strains of *S. cerevisiae*: illustration of the graph topology variation as a function of the size of $k$ (size of the sub-words used to build the compacted de Bruijn graph) using Vizitig. Higher values of $k$ tend to flatten the variations into long and unique unitigs but make the comparison of similar sequences more complex, while smaller values of $k$ depict more complex patterns on the local scope but allow identifying affiliations of sequences to genes or regions more easily.

#### Pangenomics

##### *S. cerevisiae* pangenome

To illustrate another example from the pangenome built on strains of the 142-genome study (O'Donnell et al. 2023), we focused on the *S. cerevisiae YDR018C* gene in a selection of strains in Figure S4.

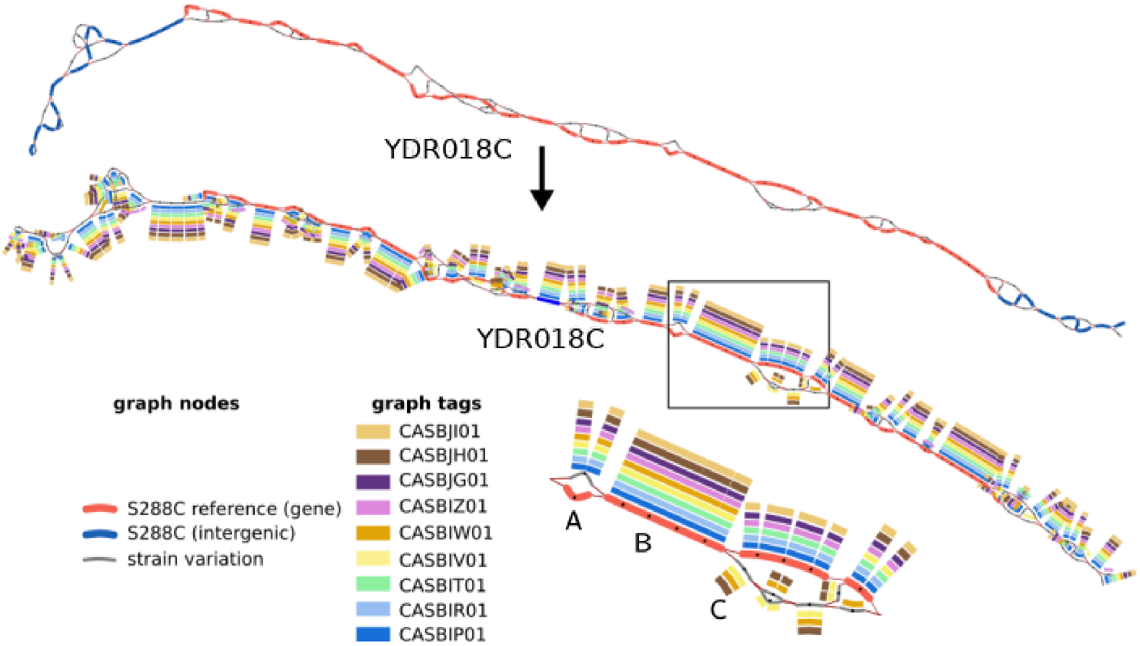


Figure S4: Close-up view of the *S. cerevisiae YDR018C* gene from a pangenome constructed using the S288C reference and nine strain assemblies. The three different graphs show the same pangenome, in three different configurations. In the top graph, *YDR018C* is shown in red, variants in grey, and intergenic regions in blue. In the middle and bottom graphs, nodes are colored by strain to highlight differences across assemblies. The square on the middle graph shows the zoomed-in region in the bottom graph. We illustrate three scenarios: **A** all strains share a variant relative to the reference, **B** all strains match the reference sequence for a given subsequence, and **C** strains CASBJI01, CASBIV01, and CASBIW01 carry variants distinct from both the reference and other strains. Variant sequences can be directly extracted for downstream analysis.

Sequences extracted from the zoomed-in region of Figure S4, for CASBIP01, CASBIW01, CASBJH01, and CASBJI01 haplotypes:

```
>61200 L: :183785:  L: :209414:
ATCTTCGAGAGGGCTCATAGGAAGTATTTTTATACTGGACT
>170545 L: :99458:  L: :183785:  L: :198673:  L: :209414:
CTGAAAAAAACATTTTCTCTAAAATTCAATATGGATCCCACCGACAGCACTAAGCTGGAAACATTTCATCTTATTGC
>198673 L: :170545:
CCACCGACAGCACTAAGCTGGAAACATTTCATCTTATTGCCTACTATACGGAAAAGGATATTCATCAAGGAAGTTTGAGGA
>209414 L: :61200:  L: :170545:
TGGGATCCATATTGAATTTTAGAGAAAATGTTTTTTTCAGCAGTCCAGTATAAAAATACTTCCTATGAGCCCTCTCGAAGA
```

#### Transcriptomics

##### Fusion events

Fusion junction coordinates were sourced from STAR-Fusion alignments provided by DepMap, using recommended filters to retain only high-confidence events. After expanding the graph from a previously identified *HM13*::*TPX2* junction, we exposed a second fusion, *BCL2L1*::*HM13*, also reported by DepMap (Figure S5A). We then applied the approach to the *CYTH1*::*EIF3H* fusion in SKBR3, identified bridging nodes between the two genes, and showed long-read support at the breakpoint, including one of the reported junctions (Figure S5B).

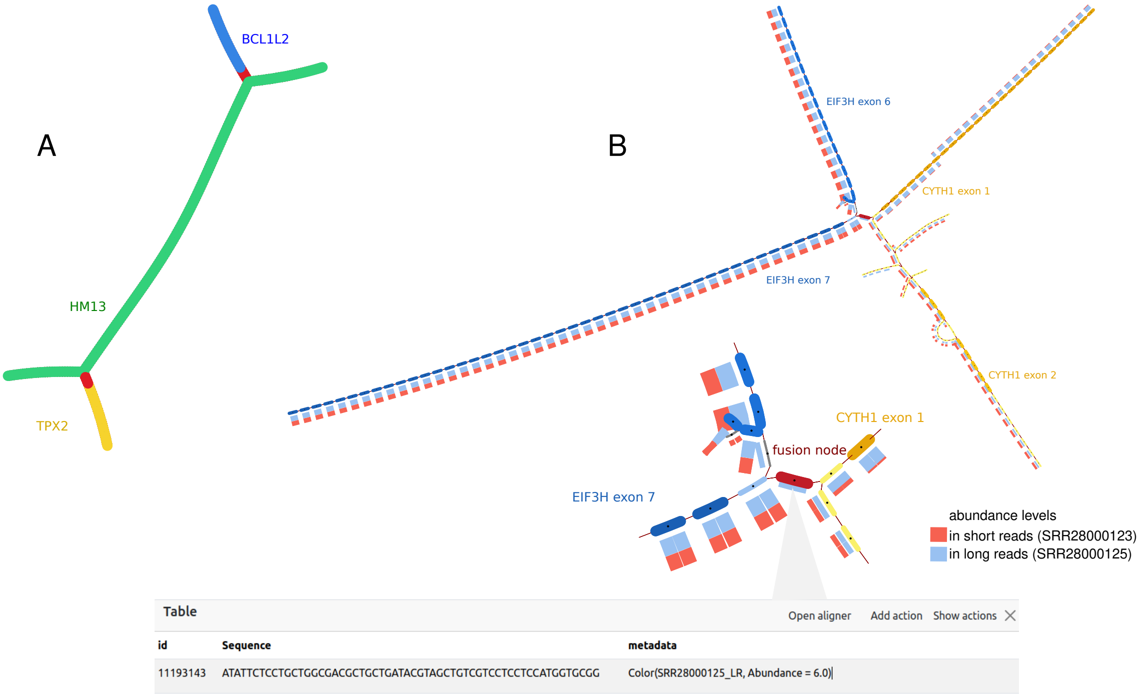


Figure S5: **A** Further exploration allows uncovering the *BCL2L1*::*HM13* fusion in sample SRR8616107. **B** *CYTH1*::*EIF3H* junction breakpoint and neighbor exons built using co-assembled short and long reads. Vizitig's sashimi-like visualization and metadata show the support of 6 long reads for the breakpoint, while short reads show no signal, as reported in the initial publication.

##### Circular RNAs

Here we extended results on the analysis of circular RNAs *MAN1A2* and *FBXW4*. Figure S6 shows additional *MAN1A2* back-splicing junctions, and Figure S7 shows the *FBXW4* circRNA.

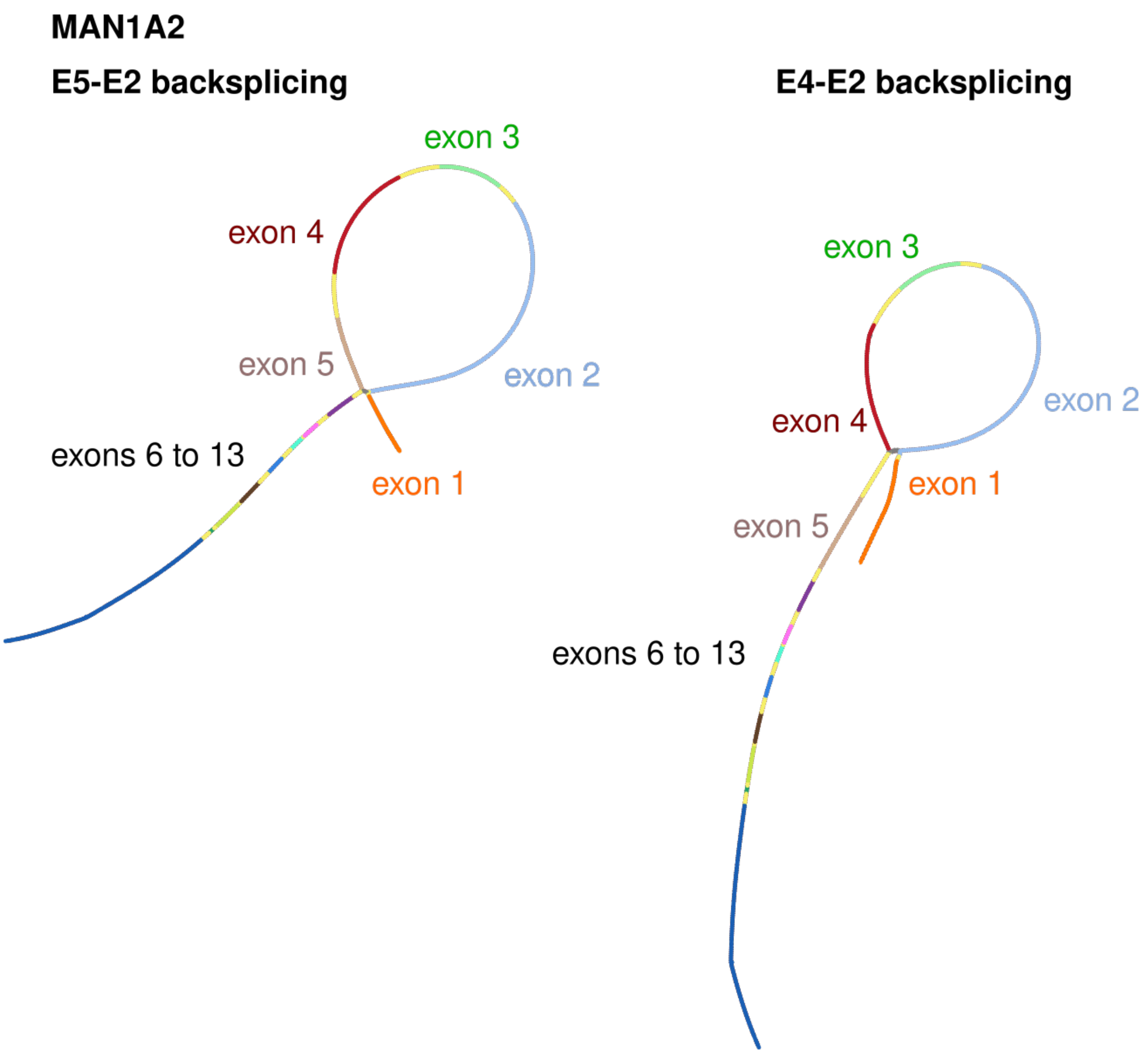


Figure S6: Back-splicing junctions in the *MAN1A2* gene supported by the literature and observed in Vizitig.

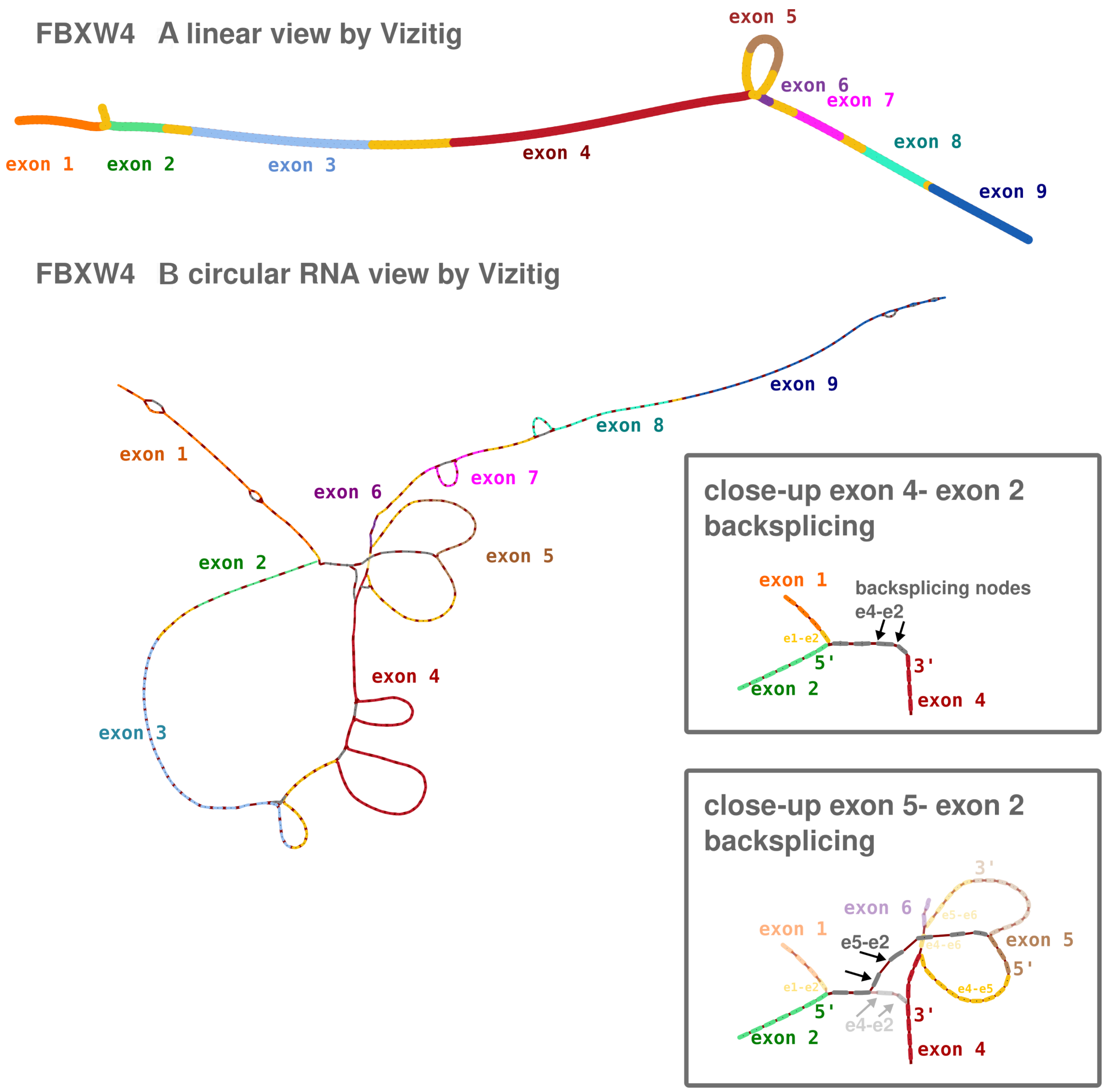


Figure S7: Different possible back-splicing junctions seen in Vizitig for the *FBXW4* gene, along with other variants (indels and alternative splicing events).

##### *C. elegans* pangenome

We built a pangenome of *C. elegans* Chromosome I using three chromosome-level assemblies from NCBI: Bristol N2 (GCF_000002985.6), Hawaiian CB4856 (GCA_000975215.1), and DL226 (GCA_022984815.1, PacBio Sequel). We built graphs with GGCAT (`-k 1023 -s 1`). Figure S8 shows 10% of the Chromosome I pangenome (right) and a zoomed-in region (left) where we illustrate different combinations of structural variants (indels) between the three strains.

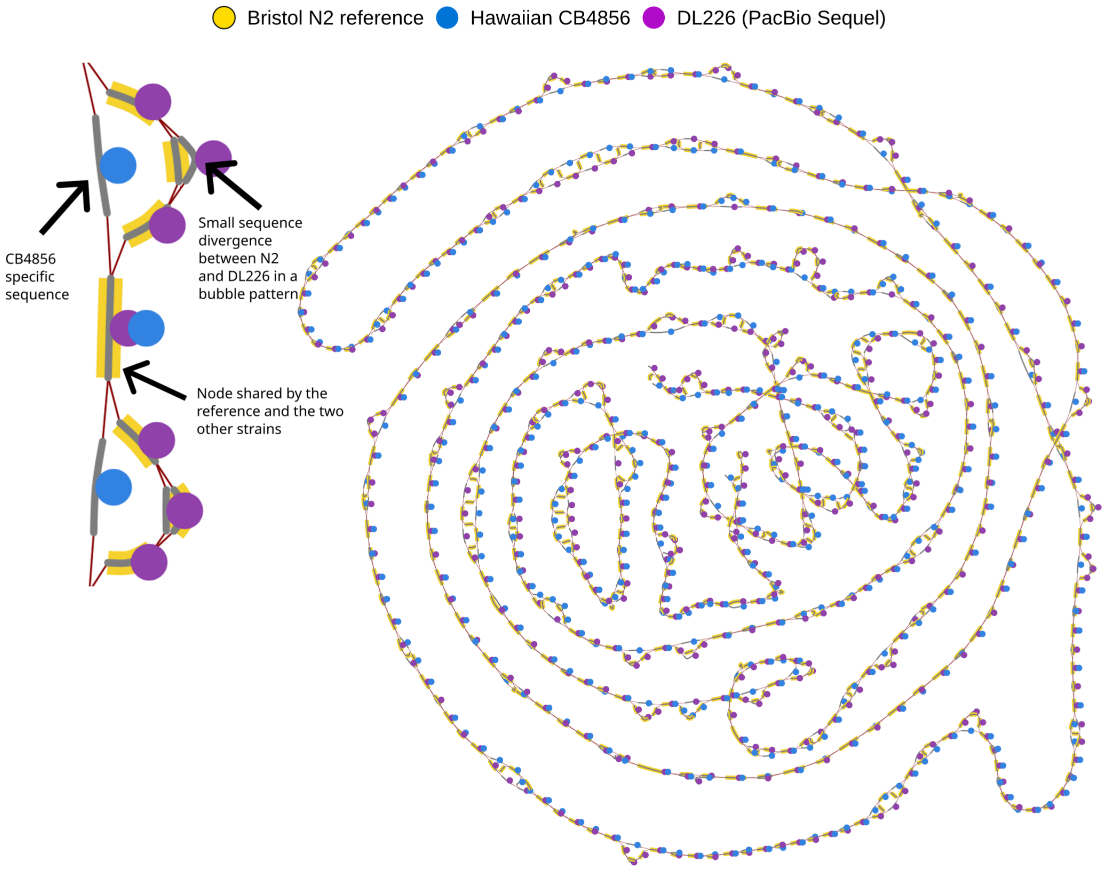


Figure S8: *C. elegans* pangenome of Chromosome I, built from three strains: Bristol N2 (yellow), Hawaiian CB4856 (blue), and DL226 (purple). Left: zoomed-in view showing shared regions (all three colors), strain-specific branches, and variants specific to each genome. Right: overview of 10% of the Chromosome I pangenome.

##### Multi-species pantranscriptome

We used Vizitig to reconstruct and visualize alternative *PAX6* (human) and *Pax6* (mouse) transcripts from transcript assemblies (`NM_000280` and `NM_001244198`) available from the NCBI, highlighting both shared, conserved exons and species-specific isoforms. By aligning exon paths across samples, Vizitig makes junction differences and sample-restricted exon usage immediately visible, helping link structural changes to potential expression differences (Figure S9).

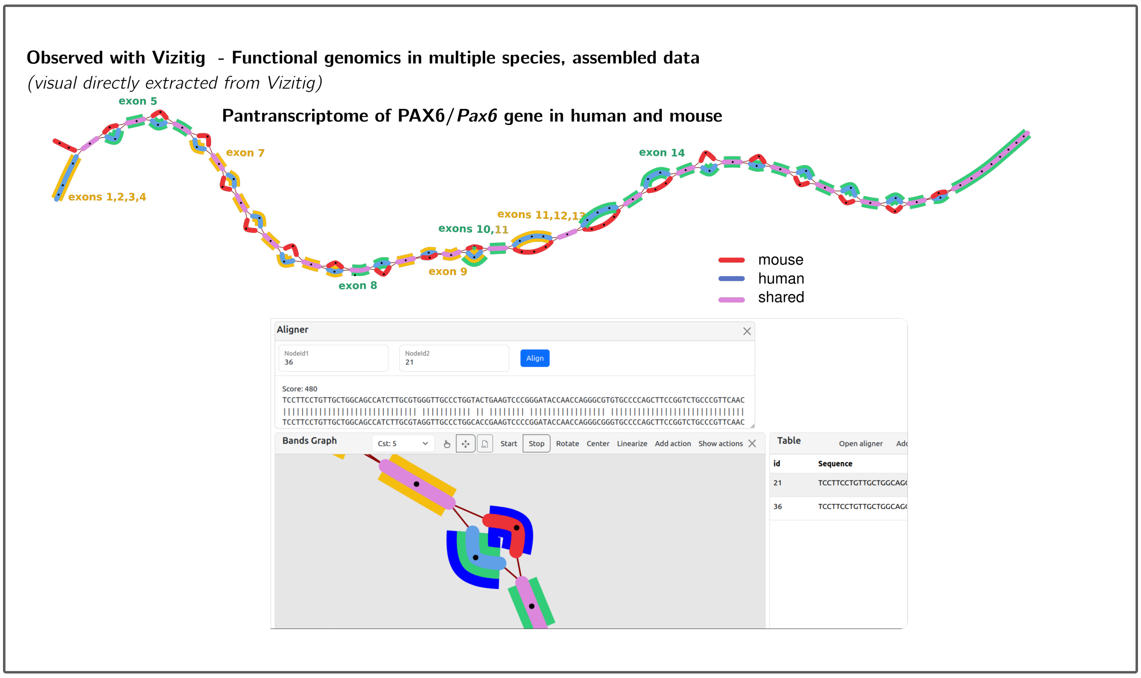


Figure S9: Top: co-assembled *PAX6*/*Pax6* genes from human and mouse and visualization in Vizitig. Red nodes are mouse sequence, blue nodes come from human, and purple nodes are shared sequences. Exons (from human annotation) alternate between yellow and green node underlines. Bottom: Vizitig's alignment tool and close-up of alternative variation in human and mouse, revealing phased SNPs in this locus.

#### Rice pangenome and *Wx* gene variants

We constructed a pangenome of *Oryza sativa* ssp. *japonica* using the Nipponbare reference genome (Kawahara et al. 2013) and 10 chromosome-level cultivar assemblies from NCBI (https://www.ncbi.nlm.nih.gov/datasets/genome/ accessions in Table S2), with $k$ =61. We annotated the *Wx* (Waxy) gene with exon boundaries and queried the locus to visualize the splice-site variant that distinguishes sticky from non-sticky rice cultivars.

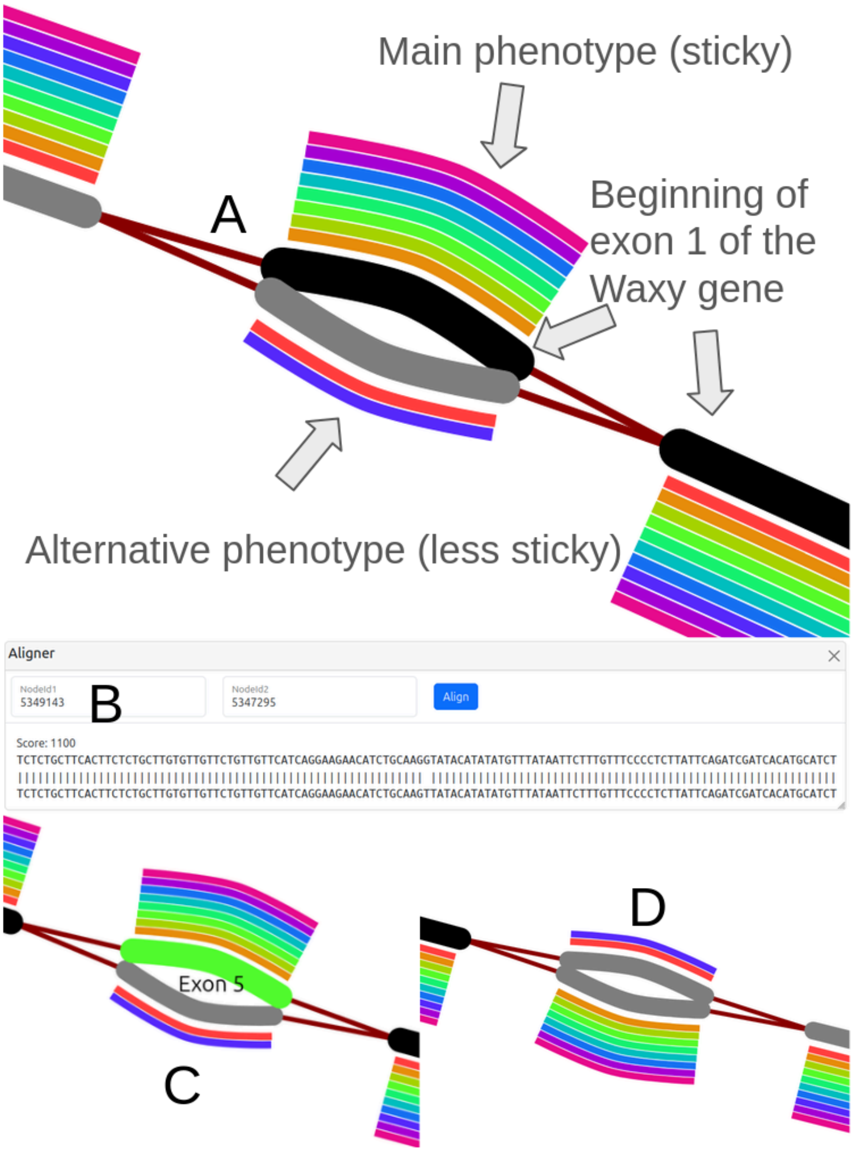


Figure S10: Visualization of the *Wx* (Waxy) gene locus in a rice pangenome of 10 japonica cultivars. **A** The beginning of exon 1 shows a branching point where sticky cultivars (main phenotype, black path) diverge from non-sticky cultivars (alternative phenotype). The G $\rightarrow$ T splice-site mutation at the exon 1/intron 1 boundary (Hirano et al. 1998) distinguishes the $Wx^b$ allele (low amylose, sticky) from the $Wx^a$ allele (high amylose, non-sticky). **B** Vizitig's built-in aligner confirms the SNP between the two paths. **C** A variant in exon 5 specific to the non-sticky phenotype. **D** An additional SNP downstream, also specific to non-sticky cultivars.

##### Splicing graphs

For RNA-seq analysis purposes, we compared Vizitig's graphs to a splicing graph library (`SplicingGraphs`, Bioconductor package in R). We compared Vizitig and splicing graph topologies on a human gene. While both topologies confirm the same variants, Vizitig gives more direct access to a sense of the sequence length and content, and a more visible rendering of the variation.

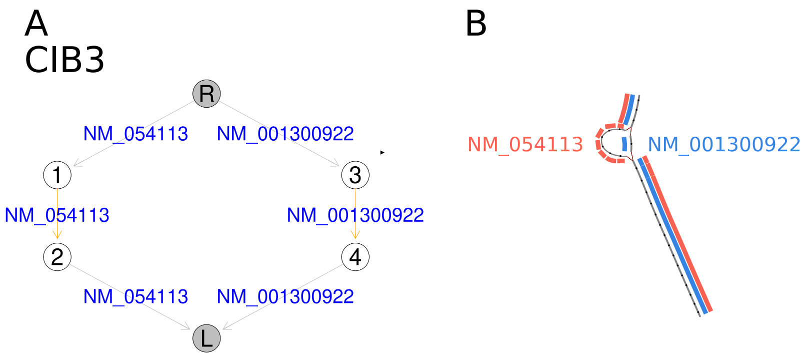


Figure S11: Comparison of the `SplicingGraphs` library in panel **A** and Vizitig graph in panel **B** to represent an RNA transcript and alternative isoforms.

#### Graph filtering

When building a de Bruijn graph from sequencing reads, the raw graph contains nodes arising from sequencing errors alongside nodes representing true biological sequences. Vizitig provides two complementary filtering mechanisms to reduce noise while preserving signal.

**Minimum $k$-mer abundance.** At build time, the user specifies a minimum abundance threshold $f$ (via the `-f` parameter). Any $k$-mer observed fewer than $f$ times across all input samples is discarded before graph construction. This removes the bulk of erroneous $k$-mers, which typically appear at low frequency, while retaining $k$-mers supported by multiple reads. For assembled genomes, $f$=1 is appropriate since assembly errors are rare. For RNA-seq data, higher values are common, depending on sequencing depth and the desired sensitivity–specificity trade-off.

**Automatic tip removal.** After abundance filtering, the graph may still contain short dead-end paths (tips) branching off from otherwise well-supported unitigs. Vizitig can automatically remove tips during visualization, as a true genomic or transcriptomic path would most of the time either continue beyond $k$ bases or loop back into the graph.

Figure S12 illustrates the combined effect of these two mechanisms on 600 RNA-seq short reads covering one gene. On row A with increasing values of $f$ (from 1 to 4, 1 meaning all $k$-mers are kept), the graph becomes progressively simpler: erroneous branches are pruned, low-covered parts of the gene are removed and break contiguity, and the remaining structure more closely reflects the spliced sequences. On row B, tip removal further cleans the graph by eliminating residual short dead-end paths.

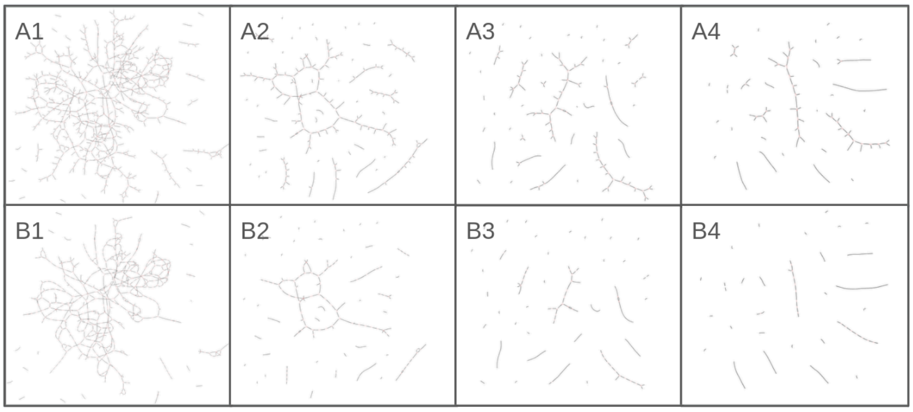


Figure S12: Effect of $k$-mer abundance filtering on graph topology. The same set of reads is used to build de Bruijn graphs with increasing minimum abundance thresholds ($f$=1, 2, 3, 4). Row A (A1–A4): as $f$ increases, erroneous branches are progressively removed, simplifying the graph while preserving the core structure supported by multiple reads. Row B (B1–B4): the same graphs after automatic tip removal.

#### Supplemental References


Garrison E, Guarracino A, Heumos S, Villani F, Bao Z, Tattini L, Hagmann J, Vorbrugg S, Marco-Sola S, Kubica C, et al. 2024. Building pangenome graphs. *Nat Methods* **21**: 2008–2012.

Hirano H-Y, Eiguchi M, Sano Y. 1998. A single base change altered the regulation of the Waxy gene at the posttranscriptional level during the domestication of rice. *Mol Biol Evol* **15**: 978–987.

Kawahara Y, de la Bastide M, Hamilton JP, Kanamori H, McCombie WR, Ouyang S, Schwartz DC, Tanaka T, Wu J, Zhou S, et al. 2013. Improvement of the Oryza sativa Nipponbare reference genome using next generation sequence and optical map data. *Rice* **6**: 4.

Li H, Feng X, Chu C. 2020. The design and construction of reference pangenome graphs with minigraph. *Genome Biol* **21**: 265.

Miao Z, Yue J-X. 2025. Interactive visualization and interpretation of pangenome graphs by linear-reference-based coordinate projection and annotation integration. *Genome Res* **35**: 296–310.

O'Donnell S, Yue J-X, Saada OA, Agier N, Caradec C, Cokelaer T, De Chiara M, Delmas S, Dutreux F, Fournier T, et al. 2023. Telomere-to-telomere assemblies of 142 strains characterize the genome structural landscape in Saccharomyces cerevisiae. *Nat Genet* **55**: 1390–1399.

Wick RR, Schultz MB, Zobel J, Holt KE. 2015. Bandage: interactive visualization of de novo genome assemblies. *Bioinformatics* **31**: 3350–3352.
